# Comprehensive evaluation of harmonization on functional brain imaging for multisite data-fusion

**DOI:** 10.1101/2022.09.22.508637

**Authors:** Yu-Wei Wang, Xiao Chen, Chao-Gan Yan

## Abstract

To embrace big-data neuroimaging, harmonization of site effect in resting-state functional magnetic resonance imaging (R-fMRI) data fusion is a fundamental challenge. Comprehensive evaluation of potentially effective harmonization strategies, particularly with specifically collected data has been rare, especially for R-fMRI metrics. Here, we comprehensively assess harmonization strategies from multiple perspectives, including efficiency, individual identification, test-retest reliability and replicability of group-level statistical results, on widely used R-fMRI metrics across multiple datasets including data obtained from the same participants scanned at several sites. For individual identifiability (i.e., whether the same subject could be identified across R-fMRI data scanned across different sites), we found that, while most methods decreased site effects, the Subsampling Maximum-mean-distance based distribution shift correction Algorithm (SMA) outperformed linear regression models, linear mixed models, ComBat series and invariant conditional variational auto-encoder. Test-retest reliability was better for SMA and adjusted ComBat series than alternatives, while SMA was superior to the latter in replicability, both in terms of Dice coefficient and the scale of brain areas showing sex differences reproducibly observed across datasets. Moreover, we examined test-retest datasets to identify the best target site features to optimize SMA identifiability and test-retest reliability. We noted that both sample size and distribution of the target site matter and introduced a heuristic target site selection formula. In addition to providing practical guidelines, this work can inform continuing improvements and innovations in harmonizing methodologies for big R-fMRI data.

## Introduction

In the past decade, brain imaging has entered the era of big data, engendering both opportunities and challenges (Charles et al., 2020; Li et al., 2019). Large-scale sample size studies enhance statistical power and can decrease false positive rates relative to single site small-sample efforts (Marek et al., 2022), making it possible to detect reliable albeit subtle differences (Button et al., 2013; Varoquaux, 2018).

Brain imaging big data roughly break down into two types, 1) single-site accumulation, represented by the Human Connectome Project (HCP, http://www.humanconnectomeproject.org/) and the UK Biobank (https://www.ukbiobank.ac.uk/); and 2) multi-site aggregation, which can be divided into prospective cohorts like Adolescence Brain Cognitive Development (ABCD, https://abcdstudy.org/) and retrospective cohorts such as Enhancing Neuro Imaging Genetics Through Meta-Analysis (ENIGMA, http://enigma.ini.usc.edu) and Depression Imaging REsearch ConsorTium (DIRECT, http://rfmri.org/REST-meta-MDD) (Chen et al., 2022b; Yan et al., 2019). The former (single-site accumulation) is costly in both time and labor required, while the latter (multi-site aggregation) inevitably entails confounders from multi-site pooling. Even the former approach is likely to encounter hardware and software upgrades, acquisition variations and other factors, which are comprised by the term “site effect” (Yan et al., 2013). Site effects can obscure neuroimaging results of interest, producing false positive effects and decreasing statistical power, undermining the credibility and generalizability of purported discoveries (Dansereau et al., 2017; Fortin et al., 2018; Yamashita et al., 2019).

Site harmonization is the process of removing or compensating for site effects, mostly through post-hoc correction. To date, the options for site harmonization of resting-state functional magnetic resonance data (R-fMRI) are few. As a non-invasive technique that allows mapping functional circuits without an explicit task (Biswal et al., 1995), R-fMRI has become a mainstream approach in the study of cognitive neuroscience and psychopathology. In 2013, Yan et al. proposed removing R-fMRI site effects by drawing on methodologies from microarray gene expression. This effort included both parametric and nonparametric standardization techniques using either additive or additive and multiplicative approaches for R-fMRI metrics, which were applied to the Functional Connectomes Project dataset (FCP, http://fcon_1000.projects.nitrc.org/index.html). This influential work underlined the importance of standardizing site effects and suggested post-hoc standardizations of mean-centering and variance-standardization. However, as R-fMRI data harmonization techniques have advanced, we undertook an up-to-date comprehensive evaluation to update recommendations on data analysis best practices and to identify needed future innovations for big-data science.

Standard evaluation procedures for a harmonization methodology should address at least two desiderata - removing site-wise heterogeneity while retaining biologically meaningful variability (Fortin et al., 2017). For the former requirement, non-significance of site statistical tests after harmonization is the generally accepted criterion, which we term the efficiency test. For the latter (retaining biological variation), various methods have been proposed. For example, Fortin et al. (Fortin et al., 2018) suggested that an increased proportion of variation explained by age as well as increased accuracy for multivariate prediction of age by harmonized measures could reflect the preservation of biological variability, thus validating the effectiveness of the approach. However, this approach still lacks ground truth of brain-wise biological representations. Considering the uncertainty of multi-voxel patterns for any kind of demographic feature, in this work we take identifiability of individuals as the objective indicator validating conservation of biological variation. Specifically, we leveraged a neuroimaging dataset containing 41 subjects who each traveled to three scanning sites (Chen et al., 2020). We assessed whether clustering accuracy (data from the same subject across different sites clustered together) of harmonized brain-wise R-fMRI metrics would be enhanced, reflecting reliable within-subject resting-state functional organization after filtering out site effects. Furthermore, this process can also support inference of whether harmonization can boost results of individual-level analyses in response to the growing need for precision medicine in which diagnosis, treatment and prognosis are tailored individually (Fisher et al., 2018; Quinlan et al., 2020).

The foundation of science is test-retest reliability, the ability of an entire experiment to be reproduced, from data acquisition to conclusions. Low statistical power can be compensated by large sample size datasets (Marek et al., 2022). Accordingly, how different harmonization methodologies affect the test-retest reliability of multicenter study is critical for big data research (Zuo and Xing, 2014; Zuo et al., 2019), but not emphasized in past evaluations. Sex as a basic biological phenotype has been investigated frequently in the R-fMRI field in relatively large-scale samples. Thus, we assess the test-retest reliability of sex differences on R-fMRI measures through quantifying overlapped statistical results by Dice coefficient between two scanning sessions within the same population compiled from multiple sites from the Consortium for Reliability and Replicability (CoRR, (Zuo et al., 2014)). Replicability refers to the ability of studies to draw consistent conclusions in response to the same research question under different experimental conditions, including population, experimental design, analyses and tools. We also evaluate the replicability of sex differences via the overlap of statistical results between CoRR and FCP, where the site effect of FCP was harmonized with CoRR. We assume that larger Dice coefficients both between sessions and between datasets indicate better data quality harmonization.

ComBat is a prevailing method for harmonizing site effects, first developed in the gene expression microarray field (Johnson et al., 2007). It decomposes site effects into additive and multiplicative components and iteratively estimates their hyperparameters with methods of moments to compute the posterior additive and multiplicative effects for each site. It is notable for its robustness to outliers, therefore it is able to handle aggregation with small sample-size datasets and is conveniently applicable. However, when estimating site effects, it does not take biological effects into account. Instead, they are simply regarded as fixed, which can be confounded with technical variation partially or wholly (Beer et al., 2020; Chen et al., 2022a; Pomponio et al., 2020). Zindler et al., reported that when using ComBat, sample distribution and usage of covariates affect the false positive rate (Zindler et al., 2020). Besides ComBat, along the data fitting spectrum various strategies aim at resolving site effects. Here we examine both parametric methods, i.e., linear regression, linear mixed regression and parametric ComBat, and nonparametric approaches, i.e., nonparametric ComBat, a subsampling maximum-mean-distance based distribution shift correction algorithm (SMA), and a representative of deep learning - invariant conditional variational autoencoder (ICVAE) (Table 1). These methods represent the mainstream of the harmonization methodology field, hence we reasoned that comparing these approaches would yield new insights for big data analyses.

**Table 1.**
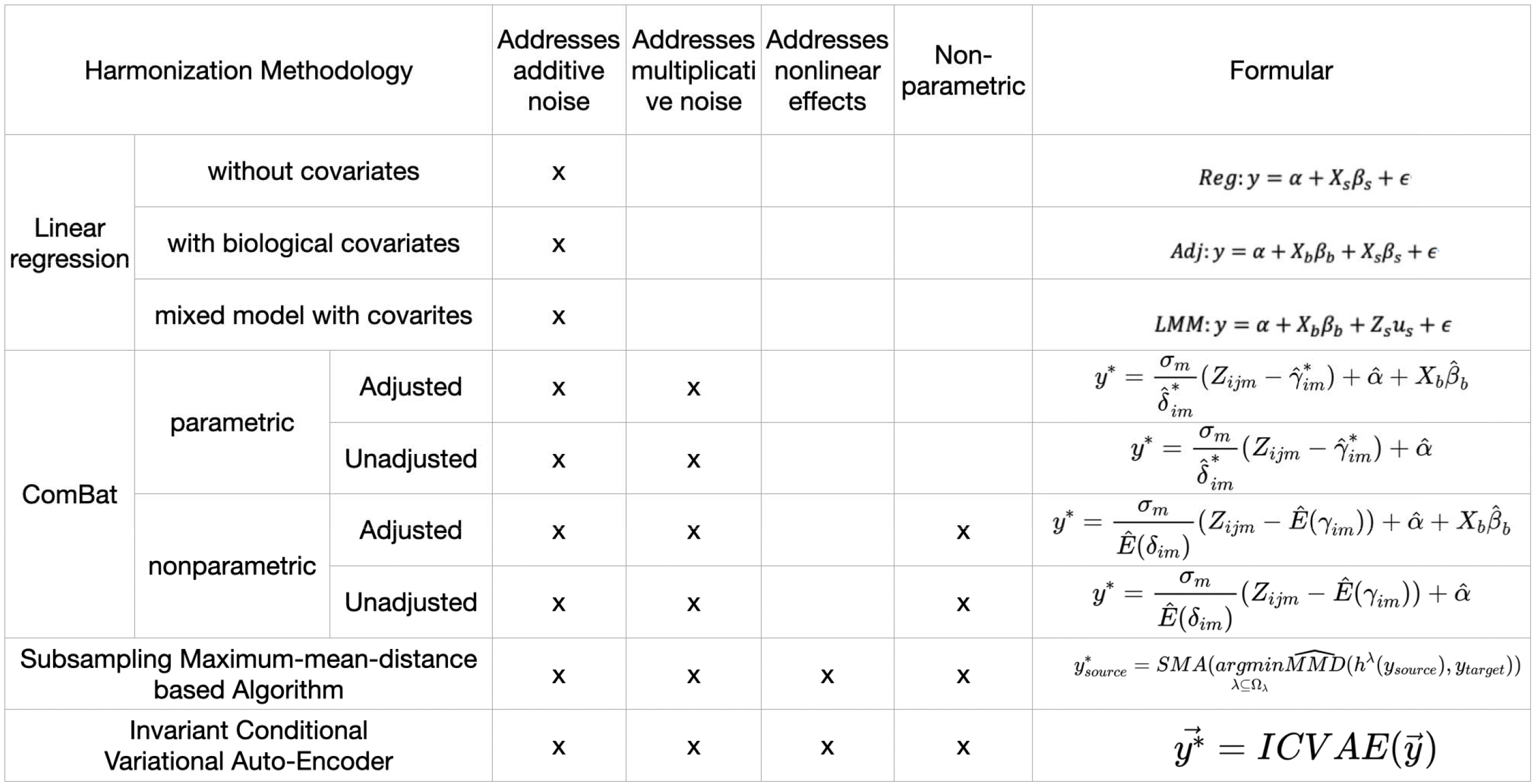
Methodology spectrum involved in the evaluation. X means yes.

We take five commonly used R-fMRI metrics, three of which reflect regional activity characteristics, namely amplitude of low frequency fluctuation (ALFF), fraction of amplitude of low frequency fluctuation (fALFF), regional homogeneity (ReHo), and intrinsic global features, namely degree centrality (DC) and functional connectivity matrix (FC), which characterize brain activity from multiple perspectives and all of which are included in the DPARSF pipeline (Yan and Zang, 2010).

In this study, we use two independent test-retest datasets and a large subset of FCP to evaluate the performance of harmonization methods (Figure 1): 1) 41 subjects traveling to three scanners (Traveling Subject Project – 3 Sites dataset, TSP-3) (Chen et al., 2020), 2) pooling sites with subjects scanned and rescanned within the same scanner, including 420 subjects from CoRR and 3) 600 subjects from 15 FCP sites. We utilize TSP-3 to investigate individual-level validity and reliability by applying the identification test, and CORR and FCP for test-retest reliability and replicability (external validity) of group-level biological effects by quantifying the Dice coefficient. First, we check basic efficiency in removing site effects through ANOVA for all three datasets and visualize harmonized results by examining P-value distributions as well as statistical output over the brain. Second, we assess individual identifiability by divisive clustering value on these measures with TSP-3 to see if the harmonization methods could preserve individual biologically meaningful information. Then we examine how harmonization approaches affect the test-retest reliability of biological differences on the same sample with test-retest sessions from CORR. Finally, we examine the extent to which biological patterns overlap between CORR and FCP after harmonization. Based on efficiency, identifiability, test-retest reliability and replicability, we conclude that SMA stands out in retaining both individual-level and group-level distinctions while eliminating site effects. Accordingly, we undertake experiments to optimize SMA to ensure best identifiability and test-retest reliability in view of the available test-retest datasets and to provide practical advice and a mechanism for target site selection.

**Figure 1.**
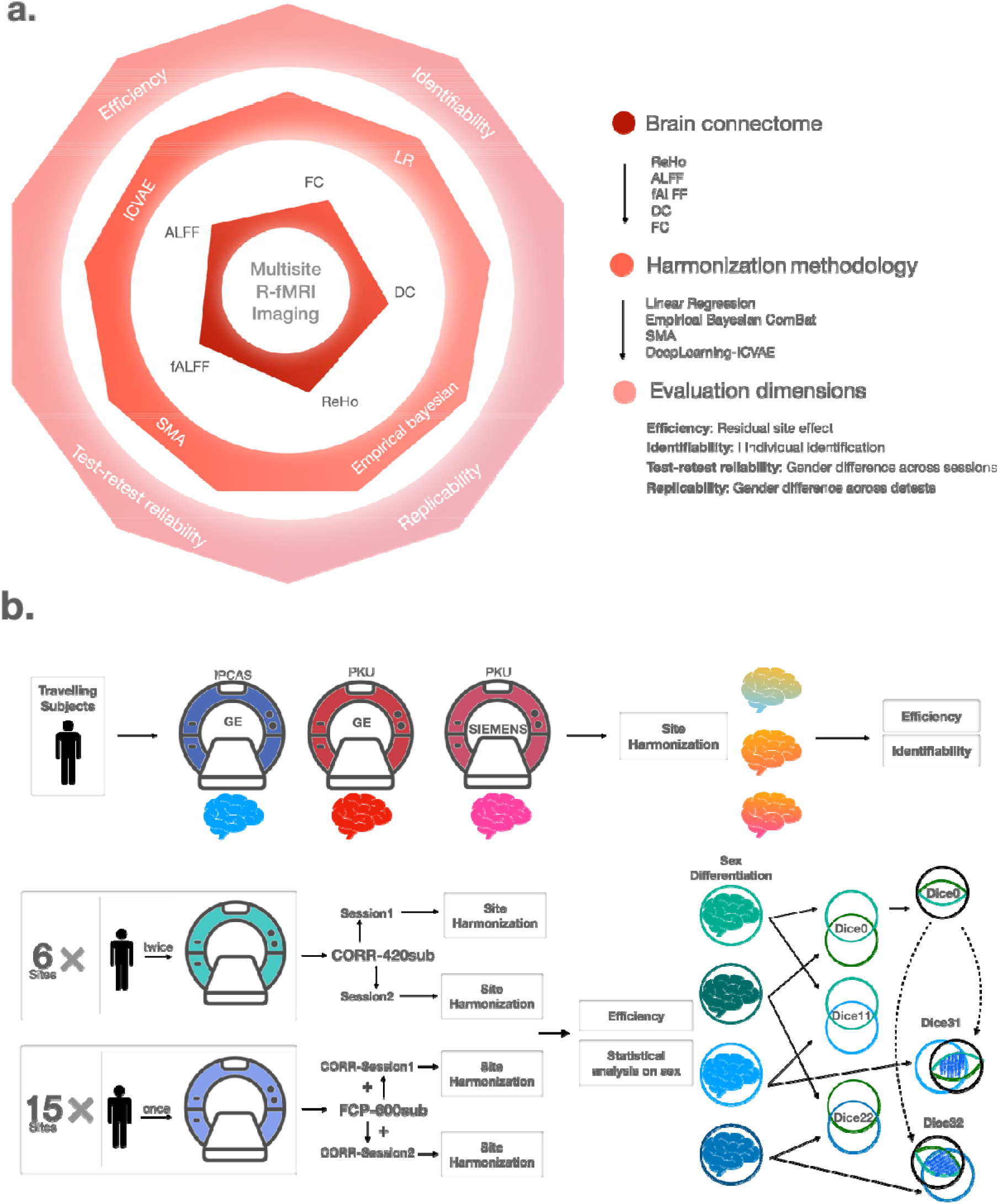
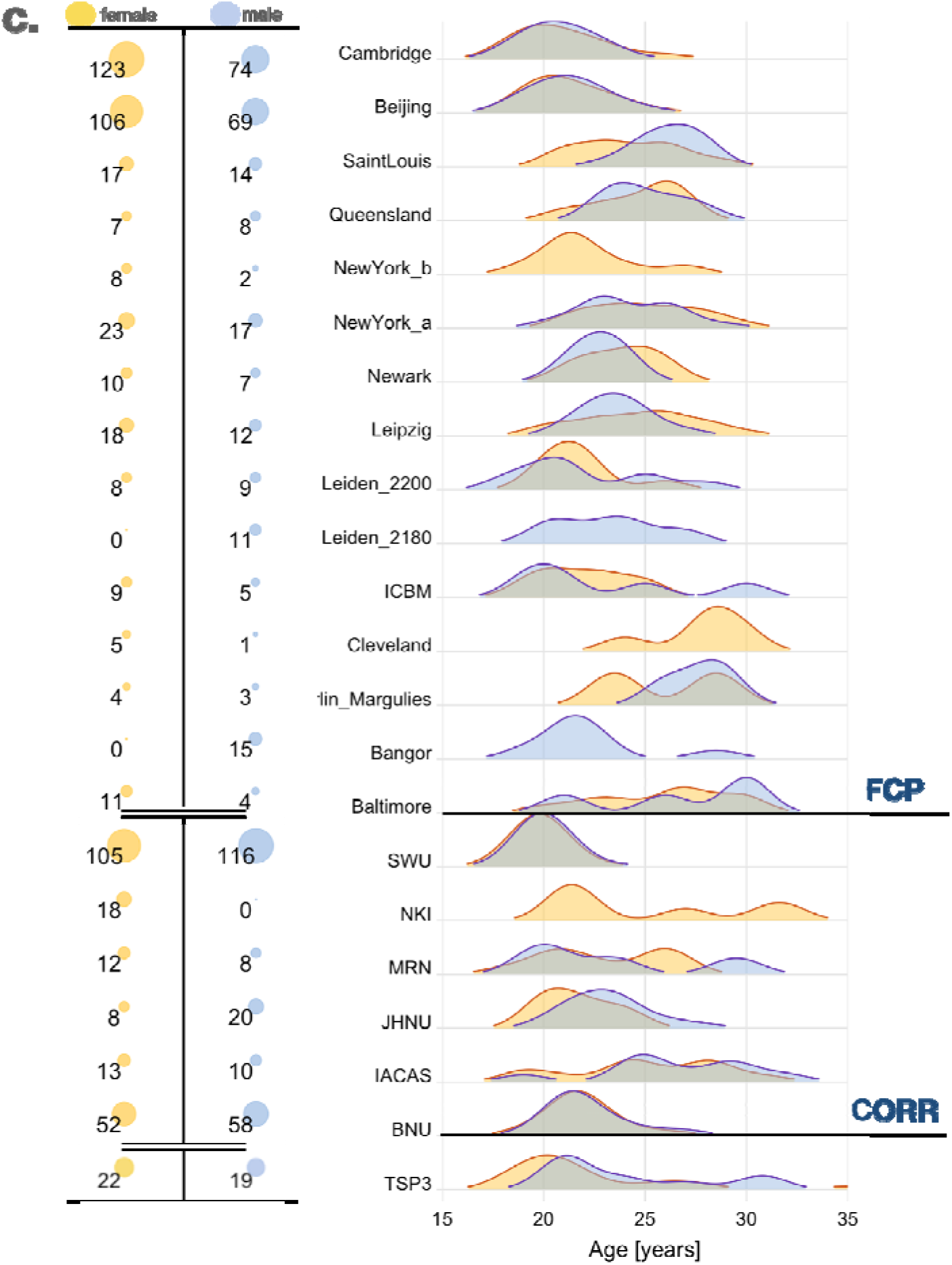
Illumination of evaluations and demographic information. **a**, The schematic of assessment framework. **b**, The evaluation process. All harmonization of site effect will be firstly tested on efficiency. Identification is run in individual-level on subjects traveling between three sites **(first row)**. Group-level statistical analysis: Dice coefficients between repeated scans of CoRR **(second row)** and across CoRR and FCP when employing site harmonization **(third row)**. It also shows the meaning of each Dice coefficient mark. **c**, Demographic information, including sites’ names, age distribution and numbers of females and males separately.

## 2. Methods

### 2.1. Datasets

TSP-3 project recruited 42 young healthy adult participants with normal or corrected to normal vision from the community. One participant who dropped out half-way was excluded, leaving a final sample of 41 (22 females; mean age = 22.7 ± 4.1 years; 40 right-handed and one left-handed) subjects. All experimental procedures were approved by the Institutional Review Board of the Institute of Psychology, Chinese Academy of Sciences, and all participants provided informed consent.

All 42 participants travelled amongst three scanners. The scan parameters for TSP-3 dataset were the recommended standardized sequence of the Association of Brain Imaging, which were developed to control site effects across different scanner models. Images were acquired on the 3 Tesla GE MR750 scanners at the Magnetic Resonance Imaging Research Center, Institute of Psychology, Chinese Academy of Sciences (henceforth IPCAS-GE) and Peking University (henceforth PKU-GE) with 8-channel head-coils. Another 3 Tesla SMENENS PRISMA scanner (henceforth PKU-SIEMENS) with an 8-channel head-coil in Peking University was also used. Before functional image acquisitions, all participants underwent a 3D T1-weighted scan first (IPCAS/PKU-GE: 192 sagittal slices, TR = 6.7 ms, TE = 2.90 ms, slice thickness/gap = 1/0mm, in-plane resolution = 256 × 256, inversion time (IT) = 450ms, FOV = 256 × 256 mm, flip angle = 7°, average = 1; PKU-SIEMENS: 192 sagittal slices, TR = 2530 ms, TE = 2.98 ms, slice thickness/gap = 1/0 mm, in-plane resolution = 256 × 224, inversion time (TI) = 1100 ms, FOV = 256 × 224 mm, flip angle = 7°, average=1). After T1 image acquisition, functional images were scanned (IPCAS/PKU-GE: 33 axial slices, TR = 2000 ms, TE = 30 ms, Flip angle = 90°, slice thickness/gap = 3.5/0.6 mm, FOV = 220 × 220 mm, matrix = 64 × 64; PKU-SIEMENS: 62 axial slices, TR = 2000 ms, TE = 30 ms, FA = 90°, thickness = 2 mm, Slice acceleration = 2, FOV = 224 × 224 mm). Multiband scan (slice acceleration = 2) was performed on PKU-SIEMENS to obtain a voxel size of 2 × 2 × 2 mm. We have shared the TSP-3 dataset online previously (http://rfmri.org/RuminationfMRIData or http://doi.org/10.57760/sciencedb.o00115.00002).

The CoRR dataset originally included 549 subjects who underwent scanning twice in the same scanner (mean interval = 205±161 days). From that dataset, 420 subjects (208 females) were selected after quality control with the following exclusion criteria. To avoid confounding of development or aging, only young adults (age between 18 to 30) were included. Subjects were excluded if their functional scans showed excessive motion (mean FD_Jenkinson ≥ 0.2 mm) (Jenkinson et al., 2002). Participants with poor T1 or functional images, low-quality normalization or inadequate brain coverage were also excluded. Further information on CoRR including scanning protocols is posted at (http://fcon_1000.projects.nitrc.org/indi/CoRR/html/index.html).

FCP is a multi-site data fusion dataset. Besides applying the same selection criteria as for the CoRR dataset, we also excluded all subjects whose scans were obtained at 1.5T. In the end, we used 600 subjects (349 females) from 15 sites. Unlike CoRR, most of the sites chosen from FCP were located in western countries, likely with more western participants, although we ignored differences in ethnicity and culture in our analyses (for details see http://fcon_1000.projects.nitrc.org/index.html). The sample sizes and age distributions by sex from each site for all datasets are shown in Figure 1.c.

### 2.2. Preprocessing

Unless otherwise stated, all pre-processing was performed using the Data Processing Assistant for Resting-State fMRI [(Yan and Zang, 2010), http://R-fMRI.org/DPARSF], which is based on Statistical Parametric Mapping (http://www.fil.ion.ucl.ac.uk/spm) and the toolbox for Data Processing & Analysis of Brain Imaging [DPABI (Yan et al., 2016), http://R-fMRI.org/DPABI]. The initial 5 volumes for TSP-3 and 10 volumes for CORR and FCP were discarded to allow data to reach equilibrium, and slice-timing correction was performed. Realignment on functional volumes were performed using a six-parameter (rigid body) linear transformation with a two-pass procedure (registered to the first image and then registered to the mean of the images after the first realignment). After realignment, individual T1-weighted MPRAGE images were co-registered to the mean functional image using a 6 degree-of-freedom linear transformation without re-sampling and then segmented into gray matter, white matter (WM), and cerebrospinal fluid (CSF). Finally, transformations from individual native space to MNI space were computed with the Diffeomorphic Anatomical Registration Through Exponentiated Lie algebra (DARTEL) tool (Ashburner, 2007).

To minimize head motion confounds, the Friston 24-parameter model (Friston et al., 1996) was used to regress out head motion effects. This model (i.e., 6 head motion parameters, 6 head motion parameters one time point before, and the 12 corresponding squared items) was chosen because higher-order models can control head motion effects better (Satterthwaite et al., 2013; Yan et al., 2013). WM and CSF signals were also included in the linear regression model to reduce respiratory and cardiac effects. Additionally, linear trends were included as a regressor to account for drifts in the blood oxygen level dependent (BOLD) signal. We did not perform global signal regression due to the ongoing controversy (Murphy and Fox, 2016). Temporal bandpass filtering (0.01-0.1Hz) was performed except for ALFF and fALFF. Of note, temporal bandpass filtering was performed after nuisance regression, so as not to reintroduce nuisance-related variation (Hallquist et al., 2013).

### 2.3. R-fMRI Metrics

ALFF (Zang et al., 2007) and fALFF (Zou et al., 2008): ALFF is the mean of amplitudes within a specific frequency domain (here, 0.01–0.1Hz) from a fast Fourier transform of a voxel’s time course. fALFF is a normalized version of ALFF and represents the relative contribution of specific oscillations to the whole detectable frequency range.

ReHo (Zang et al., 2004): ReHo is a rank-based Kendall’s coefficient of concordance that assesses the synchronization among a given voxel and its nearest neighbors’ (here, 26 voxels) time courses.

DC (Buckner et al., 2009; Zuo et al., 2012): DC is the number or sum of weights of significant connections for a voxel. Here, we calculated the weighted sum of positive correlations by requiring each connection’s correlation coefficient to exceed a threshold of r > 0.25 (Buckner et al., 2009).

To define the area for voxel-wise analyses, we first created a mask defining the overlap area for TSP-3, CoRR and FCP with a 90% group mask (i.e., the voxels selected for the group mask are included in the EPI automasks of 90% of the subjects), containing 38810 voxels. We then extracted each metric based on the unified mask for harmonization.

FC is defined as the temporal coincidence of spatially distant neurophysiological events (Friston, 2011). After we excluded cerebellum from the Dosenbach 160 atlas (Dosenbach et al., 2010), 142 cortical regions of interests (ROIs) correspond to the default mode network (DMN), sensory-motor network (SMN), dorsal attention network (DAN), ventral attention network (VAN), subcortical network (SC, instead of the limbic network), visual network (VN) and frontal-parietal network (FPN). We then computed Pearson’s coefficients for each paired ROIs’ mean time-series.

Before entering into further analyses, all voxel-wise metric maps were Z-standardized (subtracting the mean value for the entire brain from each voxel and then dividing by the entire brain standard deviation), and functional images were smoothed with a 4mm Gaussian kernel. The FC matrix was transformed by Fisher’s R to Z.

### 2.3. Harmonization Methods

#### 2.3.1. Linear regression

For linear regression series, the site factor was assumed to have an additive effect on data, to be normally distributed and orthogonal to other potential factors. We considered three methods - a linear regression model with (*Adj*) or without biological factors adjusted (*Reg*) to remove additive site effect estimated by ordinary least squares (OLS) and a linear mixed model (*LMM*) with covariates adjusted, with site as a random effect, estimated with restricted maximum likelihood (REML).

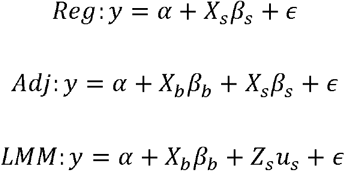

For the above formulas, the key steps included, 1) estimating the coefficients for sites (*β_s_* or *μ_s_*), and 2) subtracting estimated site effect (*X_s_β_s_* or *Z_s_μ_s_*) from y.

#### 2.3.2. ComBat

The ComBat (combating batch effects when combining Batches) method is borrowed from the technology of removing batch effects in genetics (Johnson et al., 2007). ComBat has been widely utilized to harmonize MRI data, including DTI (Fortin et al., 2017), cortical thickness (Fortin et al., 2018), and resting state functional connectivity (Yu et al., 2018). It is currently a more recognized convenient and fast site harmonization method. Compared with linear regression, ComBat further deconstructs site effects into additive (location) and multiplicative (scale) components, and then adopts the empirical Bayesian to estimate the posterior distribution to compute the mean and variance of site effects.

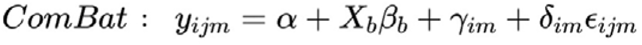

Wherein, *i* stands for site, *j* stands for individual sample index, and *m* stands for the unit of each metric (voxel or edge here). *γ_im_* and *δ_im_* represent the estimated additive and multiplicative site effects of the selected feature *m* (voxel or edge here) of site *i*, *ϵ_ijm_* is a normally distributed error term with zero mean and variance *σ_m_*.

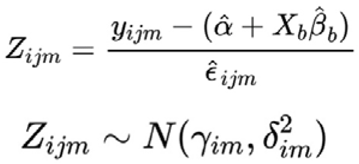

To estimate posterior distributions of standardized site effects for known samples, we deployed both the commonly used parametric method and a nonparametric approach that theoretically matches R-fMRI metrics, for which high dimensional data would be non-normally distributed.

1. For parametric ComBat, the mean and variance of a site effect hypothetically follow gaussian and inverse-gamma distributions.

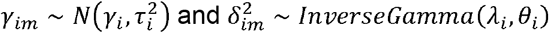 We applied the Methods of Moments to iteratively estimate means and variances until convergence.
2. For nonparametric ComBat, the prior distribution’s site-effect probability density function is undefined. Thus, we estimate the posterior expectations of the site effect parameters, denoted *E*[*γ_im_*] and 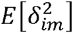, respectively, through Monte Carlo Integration.

After obtaining the site effect parameters, the empirical Bayesian was applied to estimate the conditional posterior of the site effect. And finally,

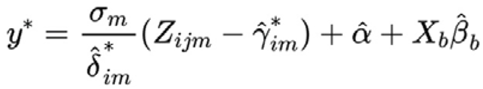

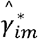 and 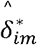 are conditional posterior site effect for each site, and the original Z value of site effect for each individual is multiplied by a normally distributed error term.

#### 2.3.3. Subsampling Maximum-mean-distance Algorithm (SMA)

SMA (Zhou et al., 2018) is an approach developed for solving statistical issues raised by data pooling, based on causal graphical model and maximum mean distance (Bareinboim and Pearl, 2016). The idea is to correct distribution shifts attributed to population characteristic differences and sample selection bias. It constitutes an integrated process from diagnosis to targeted treatment of data. Specifically, if only site-specific nuisance factors are involved, then we perform univariate direct shift correction. Extensions for fitting other co-occurring variations can be fulfilled through a subsampling framework.

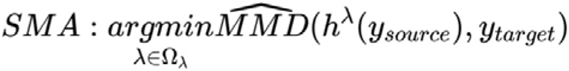

*h* here is an affine function in which *λ* represents slope and intercept choice for model selection, which are optimized towards minimization of distance between source and target distributions mapped to reproducing the Hilbert kernel space.

#### 2.3.4. Invariant Conditional Variational Auto-Encoder (ICVAE)

The idea of ICVAE is first to encode site information and the other information separately and use the encoder to learn the distribution characteristics of the original image and abstract them into a low-dimensional z vector in the intermediate hidden space. Then in the low dimensional space, the site-orthogonal information can be characterized by highly compressed orthogonal dimensions. After decoding, they can be mapped to the same dimensional space with the raw data with the same site label. The aim is to minimize the distance (defined by Kullback-Leibler divergence, KL) (Csiszar, 1975) between the generated image and the original image while minimizing the amount of mutual information between the intermediate representation z and the site information s, while controlling the encoding and decoding process to have a certain degree of correspondence. Model implementation details are shown in Supplementary Figure 1.

Moyer et al. (Moyer et al., 2020) were the first to apply ICVAE to harmonize DTI datasets. From among deep-learning approaches developed based on domain adaptation (Dinsdale et al., 2021; Zhong et al., 2020), we selected ICVAE for its interpretability.

### 2.4. Evaluation of Harmonization Effects

#### 2.4.1 Efficiency tests

To examine efficiency in removing site-wise heterogeneity, we calculated the residual site effect after harmonization with ANOVA analysis on different sites, with motion, sex and age as covariates. The two sessions of CoRR, as well as the corresponding harmonized FCP (termed FCP1 and FCP2, where numbers mark the CoRR session being harmonized) were all computed separately.

For voxel-wise R-fMRI metrics, we employed Gaussian Random Field (GRF) correction (Friston et al., 1994) on F values across whole brain with cluster p < 0.05, voxel p < 0.001, one tailed. For functional connectivity matrices, we defined 142 cortical ROIs based on Dosenbach160 atlas (Dosenbach et al., 2010) and employed ANOVA, corrected by False Discovery Rate (FDR) (Genovese et al., 2002), q=0.05.

We examined the uncorrected p distribution of site effects. In addition, we also visualized significant site effects after correction on the brain surface (Xia et al., 2013) for voxel-wise R-fMRI metrics, and a heatmap for the functional connectivity matrix showing the number of significant edges between any pair of networks based on the Yeo 7 networks (subcortical network replacing the limbic network) for each harmonization method.

#### 2.4.2. Individual identifiability

To test if the harmonization method could retain biologically meaningful individual variability, we used subject identification with divisive analysis (DIANA) (Hadi, 1991) as the clustering methodology. Through blindly clustering individual multivariate intrinsic brain representations, we obtain a square matrix in which the columns are cluster ID and rows are subject ID. For the TSP-3 cohort, 3 in each matrix element means that the three scans were sorted into the same cluster accurately. Ideally after harmonization, the matrix diagonal would contain only 3’s.

We conducted Euclidean and Pearson algorithms to construct distance matrices and chose Ward’s method (Murtagh and Legendre, 2014) as the agglomeration method, which groups data by seeking the minimum within-cluster variations summation. The accuracy formula is as follows:

*n* means the number of subjects whose repeated scans were all grouped into the same cluster and means the number of subjects

#### 2.4.3. Test-retest reliability: Dice coefficient of brain sex effect between two CoRR sessions

Test-retest reliability reflects whether a measure gives the same result for the same sample, computational steps and conditions of analysis. Here we apply harmonization steps separately for each session of CoRR, and then execute statistical analysis on sex within each session. We take GRF (voxel p=0.001, cluster p=0.05, two tailed) and FDR (q=0.05) as multiple correction methods for voxel-wise brain measures, and FDR (q=0.05) for edge-wise functional connectivity matrices.

Here we adopt the concept of Dice coefficient (Rombouts, 1998) to quantify the overlap extent between two sessions. If a harmonization strategy is better than others, it is assumed that the pattern representing sex differences will become more similar than for raw data and other approaches, with more significant overlapping results.

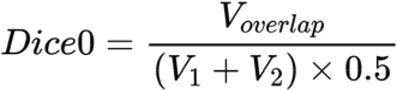

means the number of statistically significant voxels for Session1, means the number of statistically significant voxels for Session2. symbolizes the number of significant voxels spatially overlapping in both Session1 and Session2.

#### 2.4.4. Replicability: Dice coefficient of CoRR and FCP

Replicability is obtaining consistent results across studies aimed at answering the same scientific question. Here the rationale for harmonizing FCP with CoRR is to investigate whether harmonization during multi-site pooling would enhance replicability of the original data. We test replicability by calculating the Dice coefficient across different datasets after harmonization. For SMA we choose the CoRR BNU site as the target site and harmonized the FCP cohort to it. As age and sex likely affect the organization of spontaneous brain activity, based on limited studies (Dosenbach et al., 2010; van Velzen et al., 2020; Zuo et al., 2017), we took 21 years of age (>21 and ≤21) as the cutoff for R-fMRI metrics, together with sex for subgrouping. As for the other methodologies, we just combined two datasets (one session of CoRR and FCP) and took age and sex as biological covariates to regress site effects. We then performed statistical analysis on separated harmonized FCP and executed multiple comparison corrections in the same way as for test-retest reliability analysis.

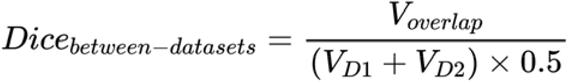

Replicability is calculated between 2 datasets, including 1) between CoRR session 1 (CoRR1) and harmonized FCP (hereafter FCP1), i.e., Dice11), 2) between CoRR session 2 (CoRR2) and harmonized FCP (hereafter FCP2), i.e., Dice22, and 3) between the intersection of CORR1 CoRR2, and FCP1 (Dice31) or FCP2 (Dice32).

### 2.5. Data/code Availability Statement

All data were obtained from publicly available online resources: TSP-3: (http://rfmri.org/RuminationfMRIData or http://doi.org/10.57760/sciencedb.o00115.00002. CoRR: http://fcon_1000.projects.nitrc.org/indi/CoRR/html/index.html. FCP: http://fcon_1000.projects.nitrc.org/index.html.

All codes used in the current study are available online: https://github.com/Chaogan-Yan/PaperScripts/tree/master/WangYW_2022. All the harmonization methods used herein will be integrated into easy-to-use modules in the next release of DPABI software (http://rfmri.org/dpabi).

## 3. Results

### 3.1. Residual site effect

For unharmonized raw data, a wide range of areas showed significant site differences after GRF correction for all datasets (Figure 2). Different site effect are observed for the same metric across datasets, possibly due to heterogeneity of scanning settings. For harmonized data, most strategies reached the goal of removing site effects, i.e., no voxels/edges showed significant site effects after multiple comparison correction.

**Figure 2.**
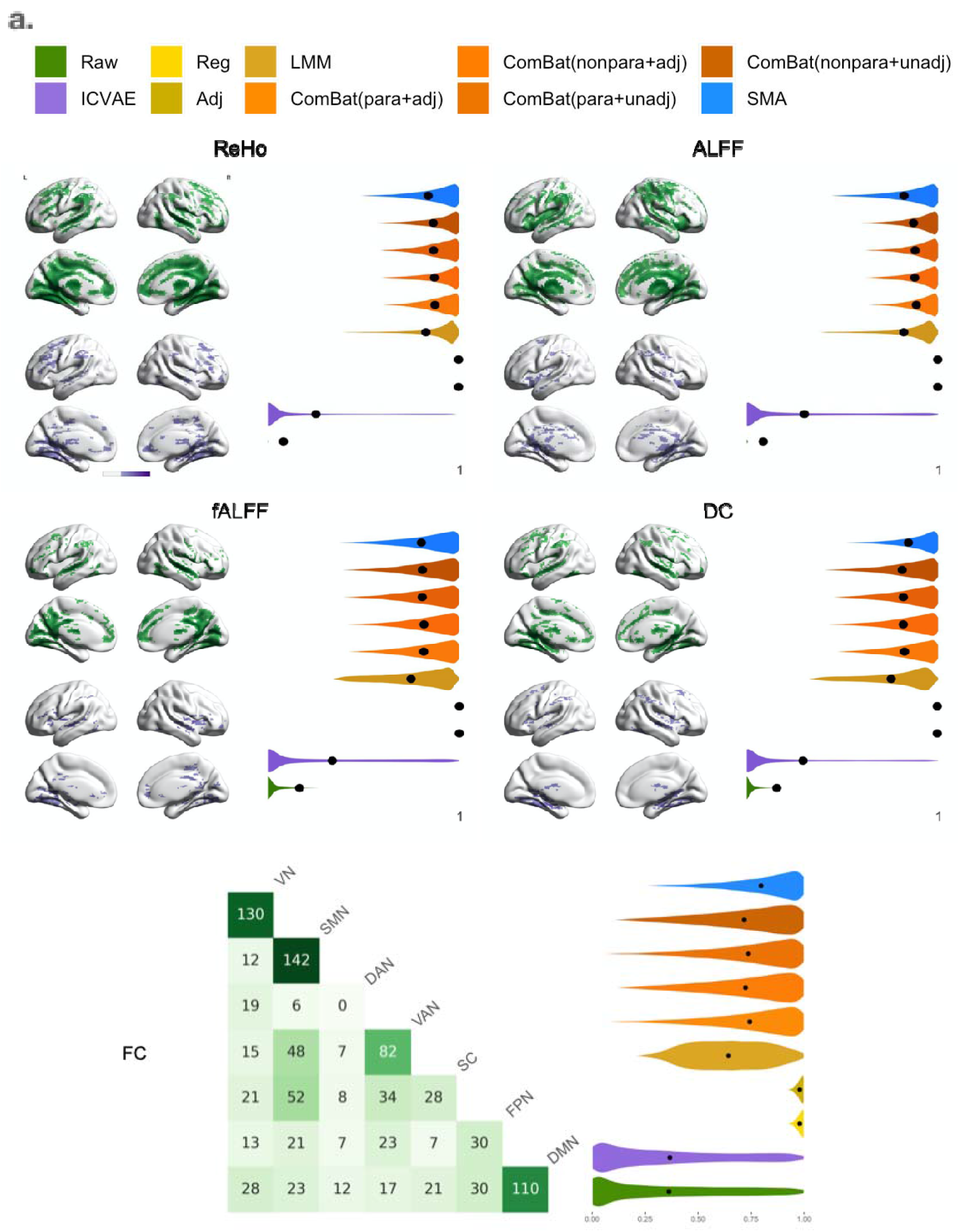
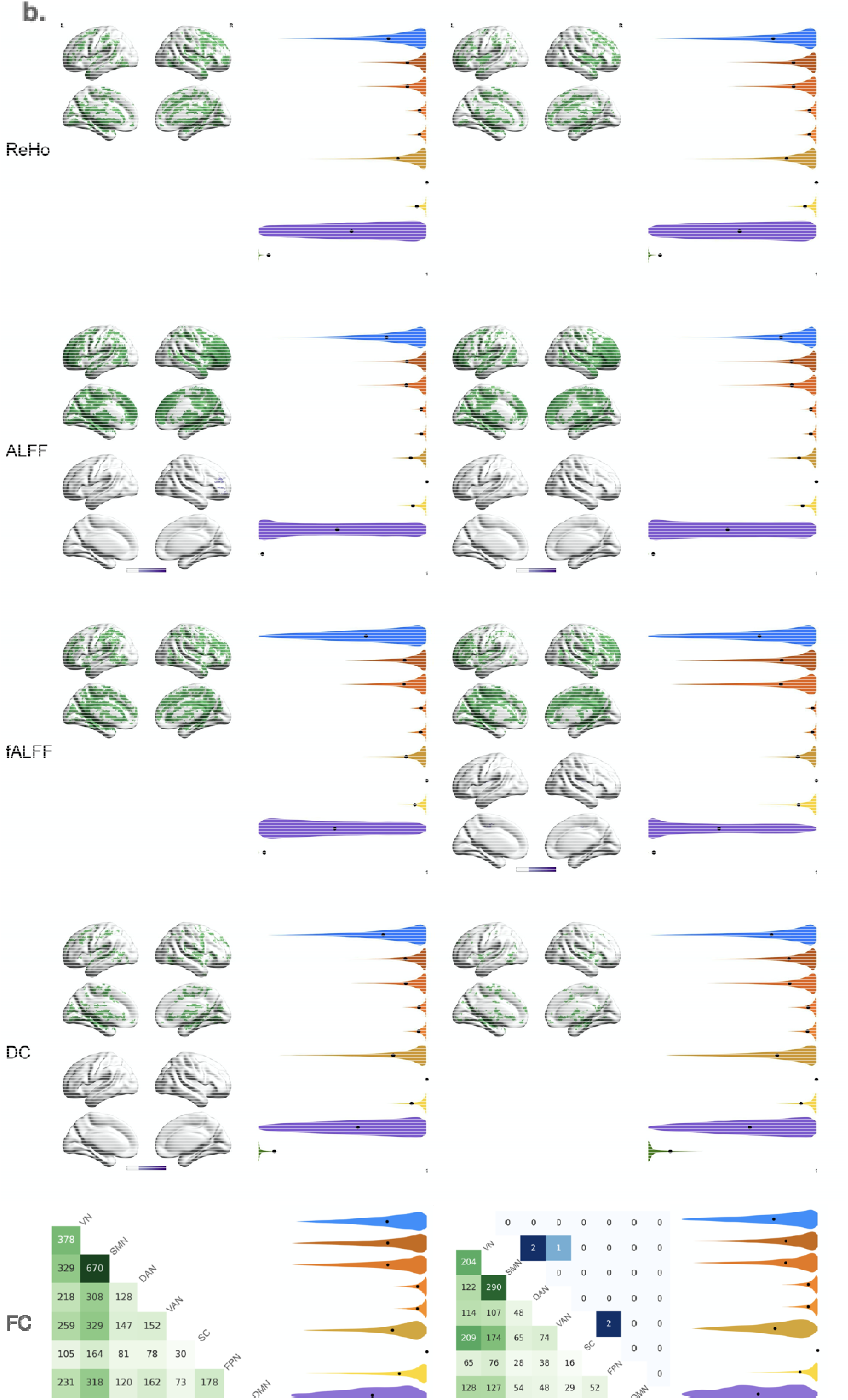
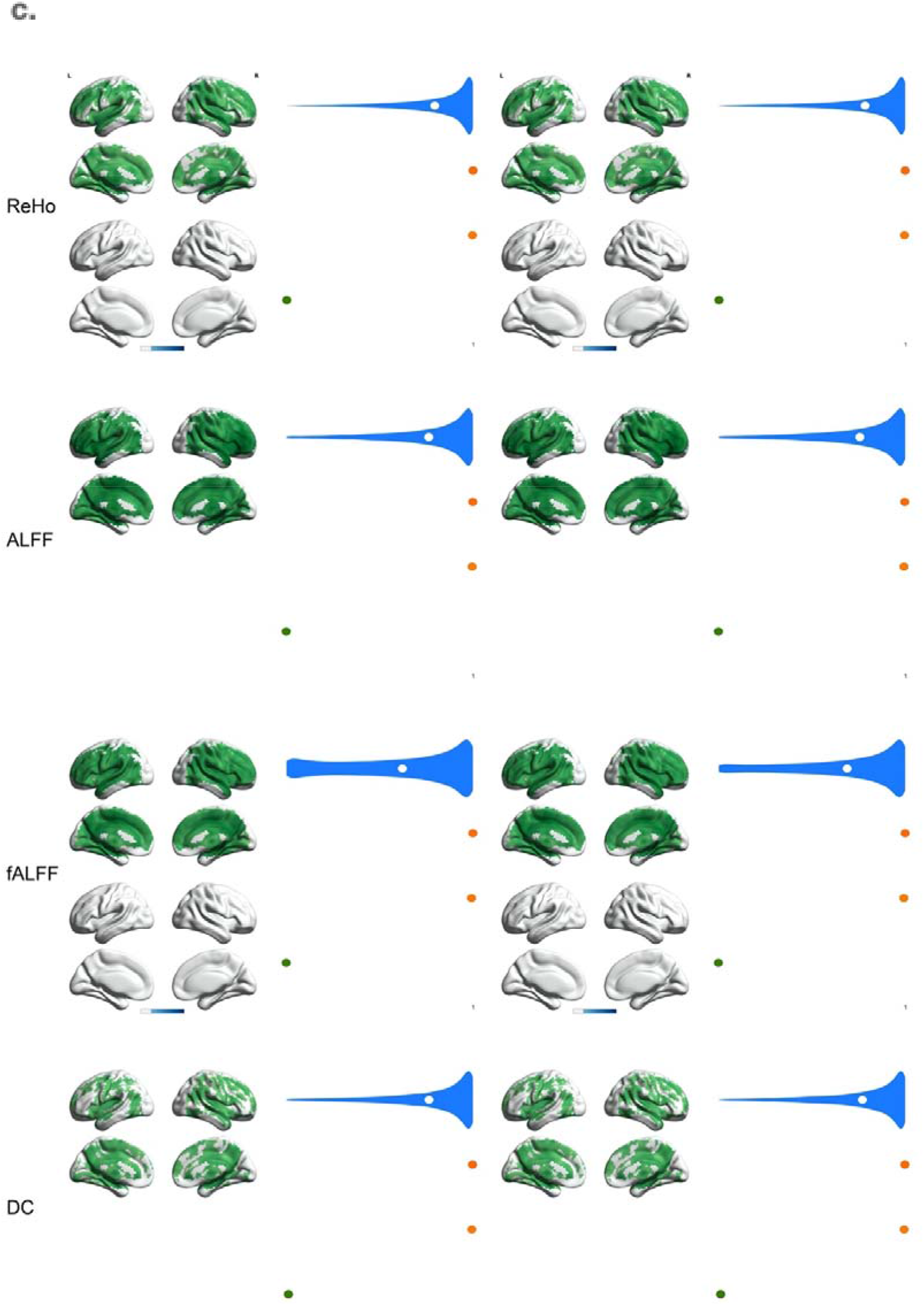
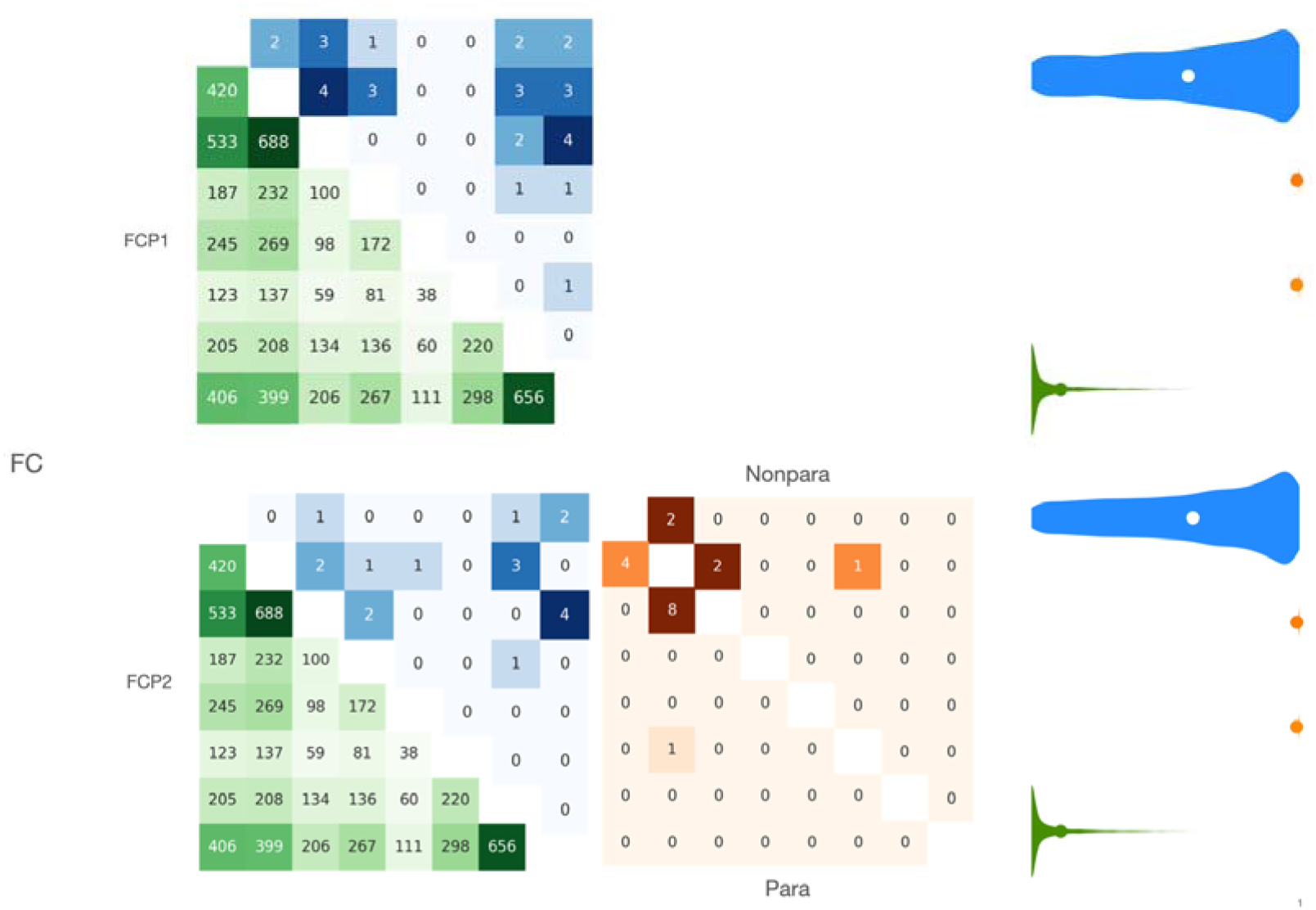
Site effects distribution across the brain. Site effect visualization for five R-fMRI metrics of **a**, TSP-3, **b**, CoRR session1 and session2, and **c**, FCP1/2 dataset accordingly. FCP1/2 represents harmonization with session1/session2 of CoRR for FCP. Different colors correspond to different methods. Colored brain area indicates significant results after GRF correction, values are log_10_(F). The darker the color, the larger the F value. Violin plots display p values of site effects. The heatmap of FC shows the number of edges with significant site effects after FDR correction between large-scale networks. The harmonization methods with no voxel/edge with significant site effect after multiple comparison correction are not presented in brain or heatmaps, whereas P-value distribution are shown for each method.

For TSP-3 (Figure 2.a), site effects consistently appeared in most area of cortex except the lateral occipital lobe in the unharmonized raw data. The largest F values converged in posterior cingulate cortex (PCC), lingual gyrus and subcortical areas such as thalamus, caudate and hippocampus. After harmonization, only ICVAE was unable to completely remove statistically significant areas. For FC, only raw data showed significant edges after FDR correction, mostly within VN, SMN, VAN and DMN. The other methods successfully removed site effects for cortical ROIs.

Significant areas in the unharmonized CoRR data were mainly found in cingulate, frontal lobe and subcortex for both sessions (Figure 2.b). From a voxel-wise perspective, site effects across the two sessions were similarly distributed. The largest F values converged in frontal lobe and lingual area. After harmonization, ICVAE basically removed site effects for ReHo and DC and little remained for ALFF and fALFF. For FC, significant site effects were mostly found within and between VN/SMN with all networks in both CoRR sessions. SMA left several edges with significant site effects in CoRR session 2.

We then performed statistical analysis on FCP1/2 to assess whether site effects had been removed. To test replicability, we only harmonized FCP with the best performing methods based on efficiency, identifiability and test-retest reliability, namely parametric adjusted ComBat (PACB), nonparametric adjusted ComBat (NACB) and SMA. From a voxel-wise perspective, unharmonized FCP site effects spanned almost the whole brain except for the occipital lobes. The largest F values converged in temporal lobes and frontal lobes. Nonetheless, significant site effects passing GRF correction in cerebellum were found after SMA harmonization for ReHo and fALFF, whereas none remained on the cortical surface for all metrics. For FC, significant site effects were mostly seen within VN, SMN and DMN, as well as between any pairs thereof. Some edges remained significant following PACB, NACB in FCP2 and SMA in both FCP1/2.

### 3.2. Identifiability

For this evaluation, we wanted to make sure that the appropriate method would remove the right amount of site effect, leaving behind enough effect size to be able to identify the same subject across multiple sites. Hence, we adapted an unsupervised machine learning technique — hierarchical clustering, to divisively cluster all 41 participants’ three brain scans (123 images in total) to see whether harmonization could improve individual clustering accuracy.

We used two kinds of methods (Euclidean and Pearson) to construct distance matrices; the detailed results are shown in Table 2. Then we ranked outputs by each distance-based accuracy calculation and averaged ranking performance of R-fMRI metrics for each method (Fig 3.a). The accuracy of clustering after SMA harmonization was the best for FC, DC and ReHo. Nonparametric-unadjusted ComBat had the best clustering accuracy in ALFF. For fALFF, except for nonparametric-unadjusted ComBat, other ComBat series and SMA had the same clustering accuracy. The pattern of Pearson-ALFF, Pearson-FC and Pearson-ReHo across brain was most individual-specific. We also noticed that: (1) the performance of the linear mixed model was worse than linear regression; (2) unadjusted ComBat outperformed adjusted ComBat in ALFF and DC; (3) ICVAE harmonized data yielded little discernible signal, with clustering accuracy for some metrics even worse than with unharmonized data; and (4) SMA retained biologically meaningful individual variability best while properly suppressing site effects.

**Table 2.**
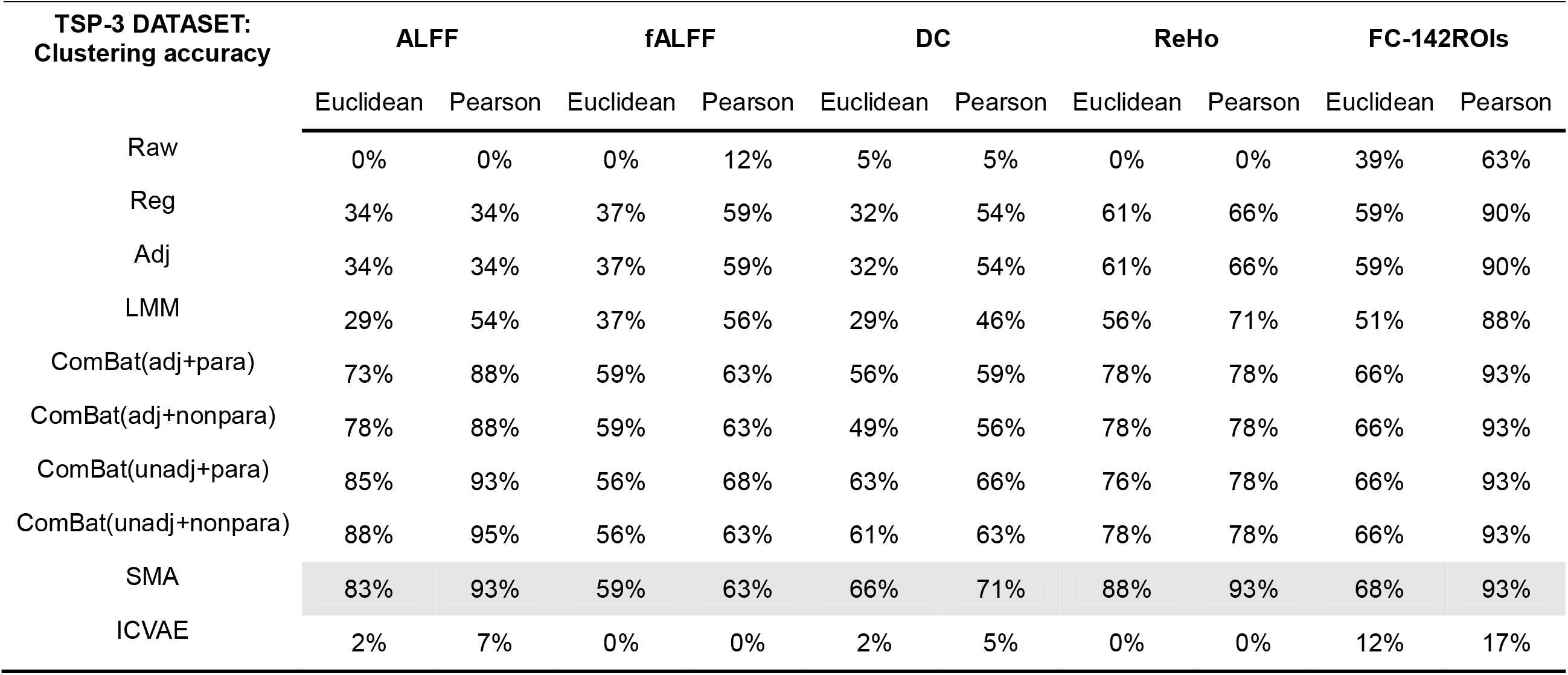
TSP-3 accuracy for each method.

**Figure 3.**
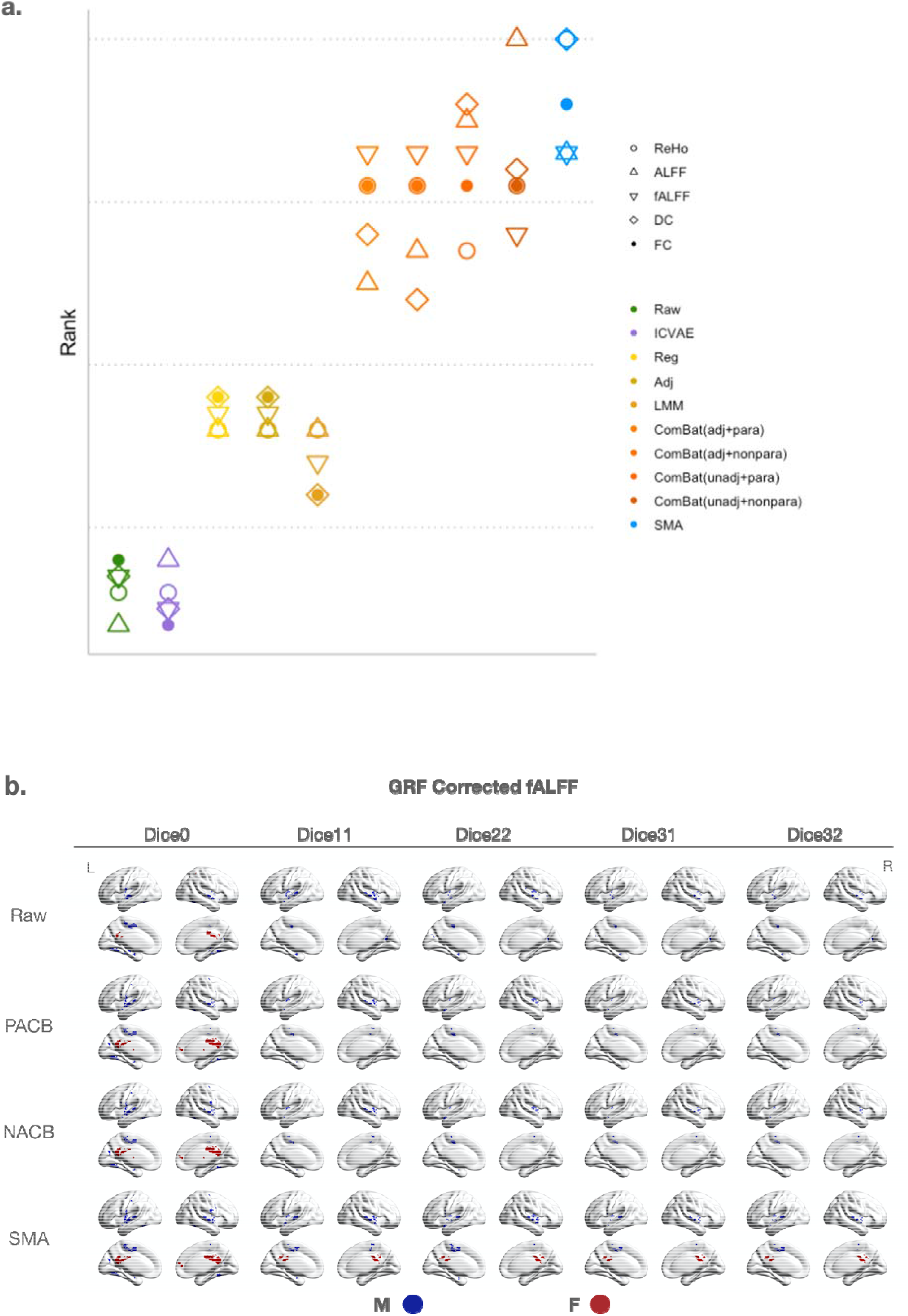
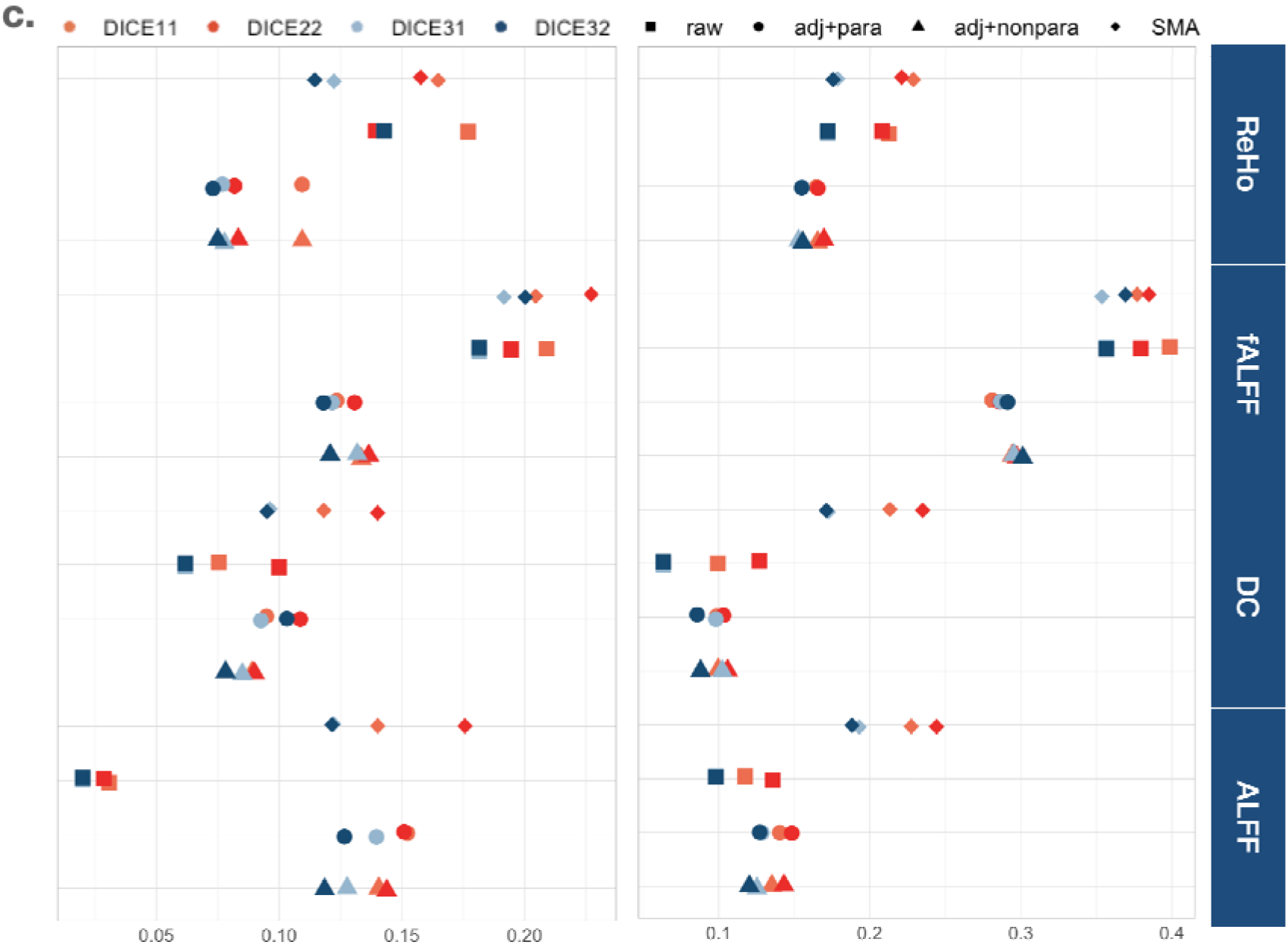
Evaluation results. **a**. Identification accuracy ranking. Colors represent methods and shapes stand for R-fMRI metric. Higher is better. **b**, Overlapped sex statistical results on brain for GRF corrected fALFF. **c**, Replicability results. Left is GRF-corrected results and right is FDR-corrected. Point shape represents methods and color distinguishes different Dice targets. CORR1 CORR2 = Dice0, CORR1 FCP1=Dice11, CORR2 FCP2=Dice22, CORR1 CORR2 FCP1=Dice31, CORR1 CORR2 FCP2=Dice32.

### 3.3. Statistical test-retest reliability and replicability

For the successful harmonization methods, we employed statistical analyses without site as a regressor to investigate the downstream impacts of harmonization.

With regard to Dice0 values, adjusted linear regression, ComBat series and SMA yielded similar overlapping extents (except that GRF corrected ReHo harmonized with unadjusted ComBat was worse). When examining the number of overlapping voxels, SMA was superior to the others in most cases. Adjusted ComBat typically yielded more overlapped voxels than unadjusted ComBat. These tendencies are presented for both GRF (Table 3) and FDR (Supplementary Table 1) corrected results. ICVAE had higher Dice0 values for fALFF and DC but the opposite for ALFF and ReHo. Considering that ICVAE and linear regression series are less appropriate for retaining biological variation based on identifiability, and that unadjusted ComBat had less area overlapping across sessions, we focused on adjusted ComBat and SMA for the following evaluation.

**Table 3.**
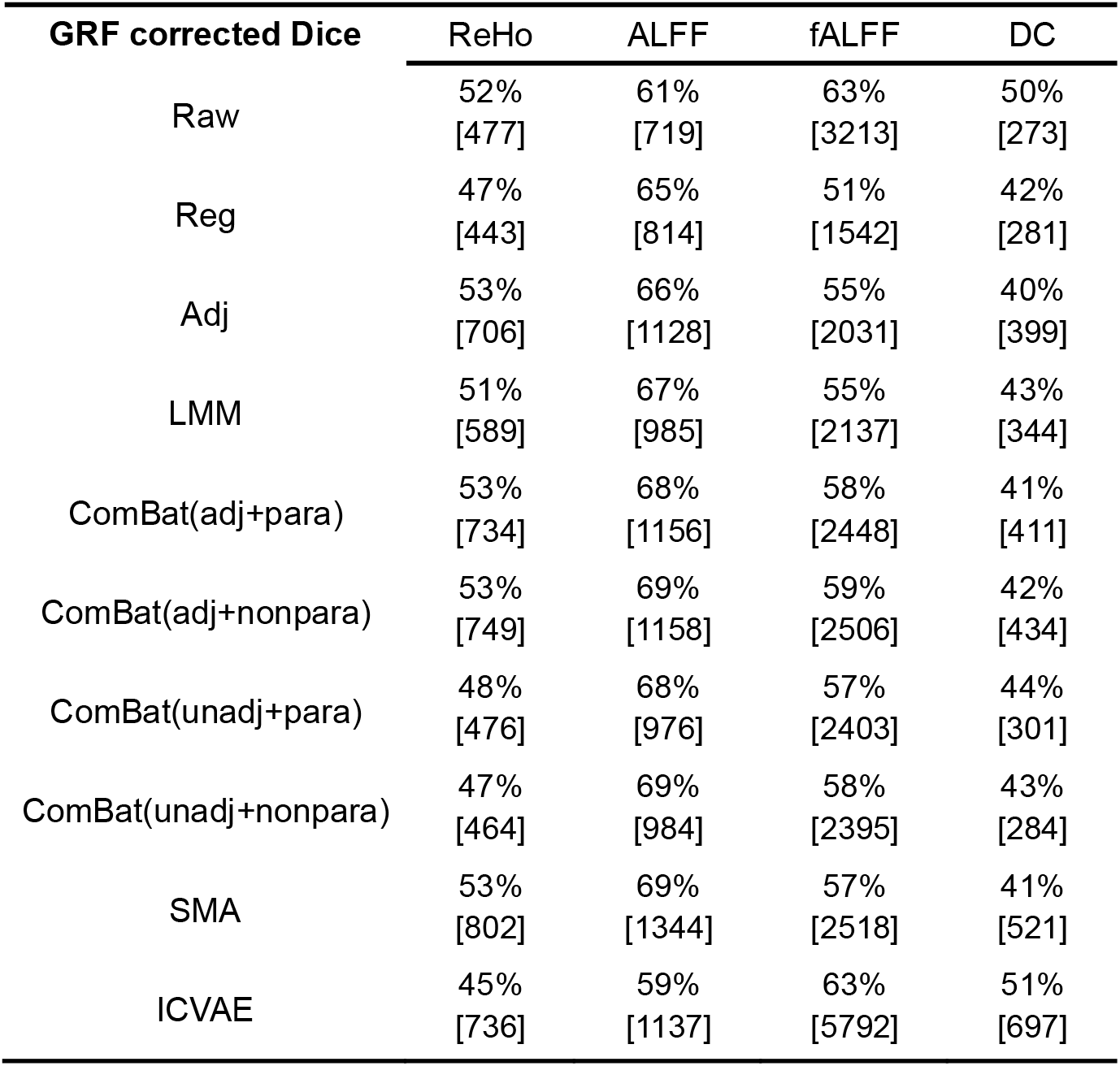
Dice coefficients for intrinsic brain metrics between two sessions of CoRR after GRF with head motion (mean FD Jenkinson), sex and age as covariates. The numbers below the percentages represent the number of voxels that were significant in both sessions.

| **GRF corrected Dice** | ReHo | ALFF | fALFF | DC |
| --- | --- | --- | --- | --- |
| Raw | 52%<br>[477] | 61%<br>[719] | 63%<br>[3213] | 50%<br>[273] |
| Reg | 47%<br>[443] | 65%<br>[814] | 51%<br>[1542] | 42%<br>[281] |
| Adj | 53%<br>[706] | 66%<br>[1128] | 55%<br>[2031] | 40%<br>[399] |
| LMM | 51%<br>[589] | 67%<br>[985] | 55%<br>[2137] | 43%<br>[344] |
| ComBat(adj+para) | 53%<br>[734] | 68%<br>[1156] | 58%<br>[2448] | 41%<br>[411] |
| ComBat(adj+nonpara) | 53%<br>[749] | 69%<br>[1158] | 59%<br>[2506] | 42%<br>[434] |
| ComBat(unadj+para) | 48%<br>[476] | 68%<br>[976] | 57%<br>[2403] | 44%<br>[301] |
| ComBat(unadj+nonpara) | 47%<br>[464] | 69%<br>[984] | 58%<br>[2395] | 43%<br>[284] |
| SMA | 53%<br>[802] | 69%<br>[1344] | 57%<br>[2518] | 41%<br>[521] |
| ICVAE | 45%<br>[736] | 59%<br>[1137] | 63%<br>[5792] | 51%<br>[697] |

We separately harmonized FCP datasets with each session of CoRR by combining the two datasets (assigning BNU as the target site for FCP in SMA). The results are presented in Supplementary Table 2 and visualized in Figure 3.c and Supplementary Figure 2. On replicability, PACB and NACB performed very similarly. SMA performed better than adjusted ComBat series by yielding more replicable results (Figure 3.b, Supplementary Figure 2). Of note, 587 subjects underwent SMA analyses, as we excluded Berlin_Margulies (all subjects were over age 23) and Cleveland (all subjects were over age 24 and only 1 was male) as their distributions deviated from the target site [BNU, only 20% subjects were over 23 years old; there should be at least 50% overlap for each subsampling factor between target and source according to Theorem 1 (Zhou et al., 2018)].

To summarize, SMA was superior for maintaining individual identifiability with higher replicability based on the above comparisons. Linear regression series and ICVAE did not perform well on individual-level identification. Unadjusted ComBat lost in detecting group-level distinctions, possibly for improper removal of site effect. Low replicability Dice values illustrate that nonparametric- and parametric-adjusted ComBat performed poorly in transporting discovered pattens across datasets. SMA was better able to harmonize R-fMRI given the above results. Given that the application of SMA still has a large number of degrees of freedom, we executed experiments based on TSP-3 and CoRR as their clean designs facilitate such investigation.

### 3.4. Optimization of SMA Target Site Choice

Comprehensively, to this point we can conclude that SMA is excellent in overall performance, particularly in identifiability and replicability. Here, we deploy experiments to figure out what kind of target site would maximize such performance.

Taking advantage of the TSP-3 dataset, we sought to identify how alteration of target site affects identifiability. We directly utilized SMA with distribution shifting but without subsampling subjects given they are the same travelling subjects. We evaluated identifiability fluctuations depending on target site selected; accuracy results are shown in Supplementary Table 3. As shown by Fig 4.a, rearranging target sites barely produced differences, although fALFF and DC were more sensitive to variation of mechanical properties.

**Figure 4.**
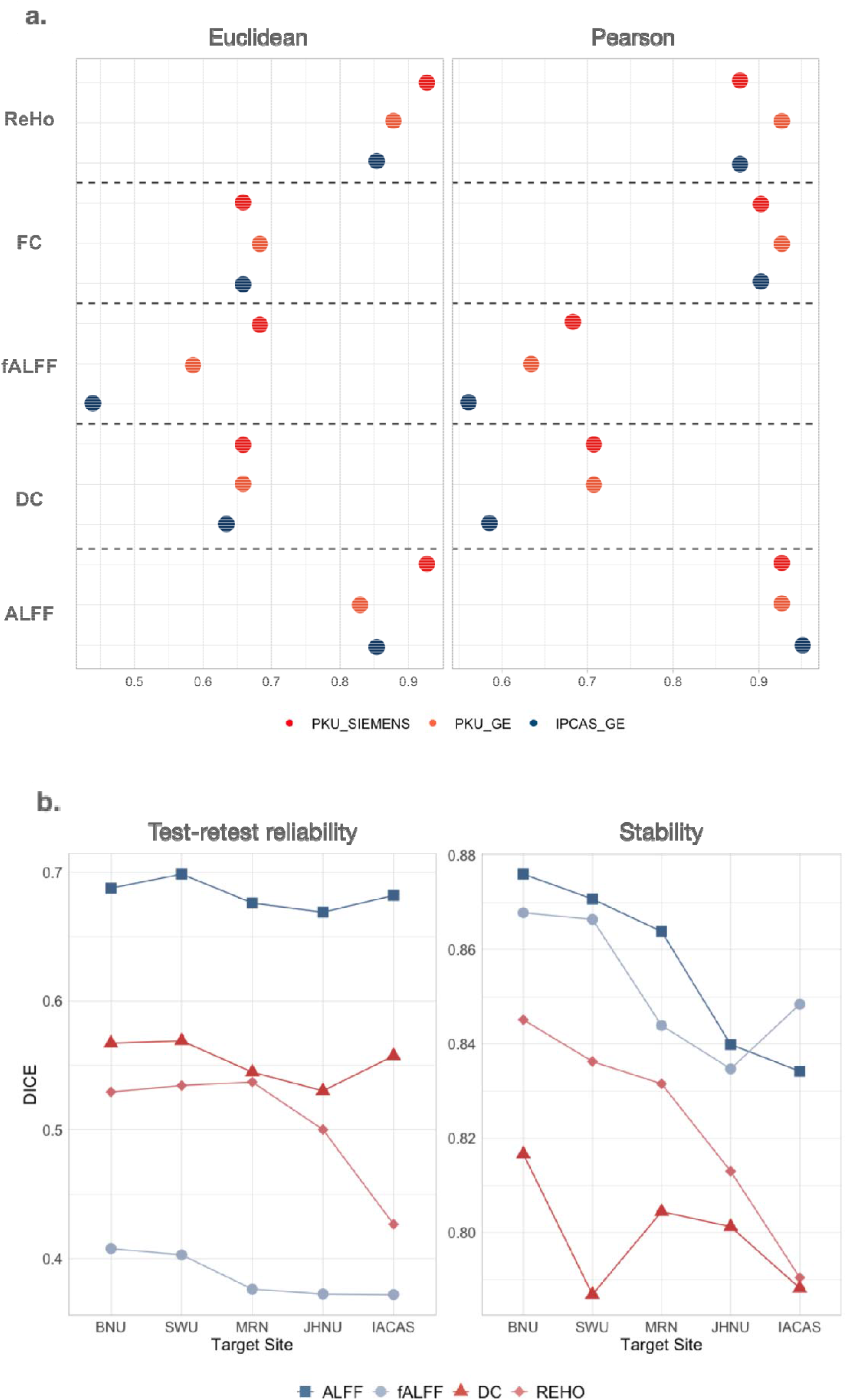
Site choice experiments. **a**, TSP-3: Identifiability accuracy when choosing different sites as the target. Colors correspond to sites **b**, CoRR: Test-retest reliability and stability when choosing different sites as the target. The x-axis is arranged in order of site selection formula, left to right best to worst. Color and shape of points distinguish R-fMRI metrics. Only GRF correction is displayed.

To investigate if target site choice would change test-retest reliability, we varied CoRR target sites and compared the mean Dice coefficients. Specifically, we first employed SMA with different target sites, following the same subgrouping rules. Then we examined sex differences by T-tests on the harmonized results, applied the same correction method and threshold settings, and calculated the Dice coefficients for the two test-retest sessions (test-retest reliability). As displayed in the left part of Figure 4.b and Supplementary Figure 3, test-retest reliability fluctuated depending on target site choices and R-fMRI metrics. ALFF, fALFF and DC changed unanimously across distinct target sites: higher Dice values were found at large-sample sites. For ReHo, test-retest reliability increased to the maximum at MRN and then turned sharply down. Overall, larger sample sizes were associated with higher test-retest reliability (BNU 110 subjects, SWU 211 subjects).

We also tested if the results of one target site were more stable than other sites, i.e., if the results are similar to those using other sites as target. For a given target site, we calculated the overlap (Dice coefficient) with results when setting each of the other sites as the target site. We then averaged the Dice values across the pairs and two sessions for each site [i.e., mean of 8 Dice values, e.g., BNU1 ⋂ SWU1, BNU1 ⋂ MRN1, BNU1 ⋂ JHNU1, BNU1 ⋂ IACAS1, BNU2 ⋂ SWU2, BNU2 ⋂ MRN2, BNU2 ⋂ JHNU2, BNU2 ⋂ IACAS2 (BNU stands for the target site abbreviation, and 1/2 for the harmonized session of CoRR.)]. As Figure 4.b right, Supplementary Figure 3 right and Supplementary Table 4 show a clear descent across all R-fMRI metrics, starting from BNU and ending at IACAS (exceptions of fALFF at JHNU and DC at SWU). While SWU had a larger sample than BNU, the age range was narrower; the age distribution of IACAS offsets to late 20s and was distinct from other sites. In that case, we further probed the distances between samples of target site and source sites. Specifically, we first subgrouped samples according to population characteristic differences and selection bias specific to our case (classifying by sex and 21 years of age) to compute the percentages for each subgroup, which is a rough depiction of a site’s demography. Then we quantified the discrepancy of demographic distributions through Kullback-Leibler (KL) Divergence. Here we hypothesized that if a site is a better target, its demography would need the fewest extra samples to reproduce the others (i.e., source sites). Therefore, we ranked the average value of KL distances between a site and those left out. Site MRN needed the fewest extra samples regardless of the sample number (Mean KL value: MRN<BNU<JHNU<IACAS<SWU<NKI).

These phenomena indicate that both sample size and subgrouping factors’ distributions between source sites and the target site modulate harmonization and downstream analysis. The larger the sample size of a target, the better the test-retest reliability. It is also beneficial that the subgrouping factor distribution of a target site be as similar to that of as many source sites as possible. Given the above experience we provide a heuristic formula for target site selection:

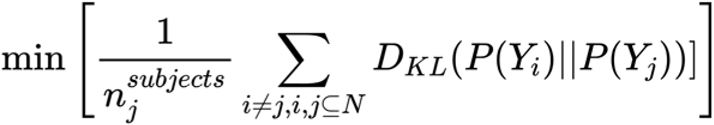

*Y* represents subgroup labels, *j* for target site ID, and *i* for source site ID. and stand for subgroup percentages in source site and target site. refers to Kullback-Leibler Divergence calculator. Here, it quantifies the amount of extra information for site j needed to characterize the population differences and sample selection of site i, based on the corresponding sample features of site j. N means the set of site IDs and means the number of subjects in site j. Finding the minimum value is equivalent to finding which site needs the least extra information to encode all other sites in terms of the relevant demographic factors, while prioritizing relatively large sample size. Here we do not consider data quality.

Based on the above tests, we give our application advice as follows:

1. **Site chosen**: The ideal situation is large sample-size sites with relatively even distributions, which means sample numbers in each division are comparable to those of source sites, thus reducing bias induced by sample selection. If such a site does not exist, we prefer the largest sample-size site containing enough demographic range (such as BNU or SWU) in both continuous and discrete biological factors. Suboptimal choices are those with moderate sample size but even demographic distribution (such as MRN). We provide an empirical target selection formula for reference.
2. **Subsampling**: This mainly depends on the hypothesized relationships among factors. Zhou et al., (Zhou et al., 2018) used a graphical causal model to depict the causal relationships among age, diagnosis and other endogenous variables influencing CSF measurements. In our example, we regard both sex and age as impacting R-fMRI metrics and demographic characteristic difference *E_P_* and selection bias *E_B_* directly affecting age and sex. Here we take 21 years old as the age division threshold based on existing evidence.

> **Briefly, a site with large sample size, whose distribution is uniform or similar to the population one intends to make conclusions about, would be an ideal target site.**

## 4. Discussion

To evaluate appropriate brain imaging harmonization methods for the big data era, we started by simplifying population composition by examining the same participants travelling to three scanners. When evaluating identifiability, SMA was the top performer with superb control of bias and variances. In the next step, we examined test-retest reliability of biological differences in multisite large sample datasets with each participant scanned twice on one scanner. SMA produced the largest number of overlapping significant voxels while having comparable Dice coefficients to ComBat, potentially indicating sensitivity to sample-specific nuances. Finally, generalizability across datasets was apparently enhanced by SMA. Among such results, SMA-harmonized detected higher activity of posterior cingulate cortex of females for both ALFF and fALFF, while ComBat-harmonized was unable to transport that discovery from CoRR to FCP. Significantly higher fALFF in males was found in bilateral insula, sensory and motor related cortices, consistent with prior studies (Al Zoubi et al., 2020; Allen et al., 2011). In addition, areas with significant fALFF were highly spatially similar to those reported by Biswal et al. (Biswal et al., 2010), showing high replicability and statistical power after SMA harmonization. By leveraging TSP-3 and CoRR, we sought how to best select a target site to optimize harmonization results. Assessments of identification and test-retest reliability show that the site with the largest sample size and widest overlap distribution with all other sites should be selected as the target site.

We noticed that the ComBat series have decent efficiency and impressive test-retest reliability performance. Here we try to elucidate the reason for their unsatisfactory performance on our other indices. The ComBat algorithm can be decomposed into a linear part, which has the same logic as normal linear regression, and the standard residual variance part, which normalizes each site’s error variance by dividing by the Bayesian-estimated site-wise standard error. The ComBat model can only include linear relationships between independent and dependent factors. In our case, the linear hypothesis apparently impaired identifiability yet enhanced test-retest reliability, since dependence between estimates of slope and class value could lead to an artificially increased class signal. Moreover, the way ComBat standardizes sites with once-for-all estimation of site distribution hyperparameters would diminish the contribution of small-scale sites. The more scanners are involved, especially when most sites have no more than 30 subjects, the larger is the estimated standard deviation of site variance. As a consequence, biological diversity among groups could be diminished, which results in insensitivity to moderate contrasts, reflected in 1) fewer significant voxels compared with SMA with the same number of subjects and 2) large test-retest reliability but lower replicability with some significant areas missed.

SMA was most able to balance individual-level differences, exemplified by high accuracy in identification clustering, and group-level characteristics, illustrated by stably higher Dice coefficients in both test-retest reliability and replicability. These quantified qualities confirm SMA’s preeminent role for R-fMRI site effect harmonization. We further elaborate on SMA’s highlights in harmonization. On the whole, SMA harmonizes site effects through one-on-one “coaching” (the target site “teaches” the source sites), which prevents biasing toward large-sample sites. Meanwhile, its subsampling algorithm retains enough information of interest and offsets the influence of outliers and sample size. Zooming in on the technical details, the subsampling scheme has two advantages. One relates to “landmark” data (Gong et al., 2013). The landmarks in the harmonization condition are those matched subgroups coordinated by Z variates, corresponding to d-separation variables in a graphical causal model. This trick mainly controls the matchiness of biological and other covariates between sites and keeps features from the source site adaptive to the target site. The other advantage is resampling, which can make the results robust to outliers even with small sample sizes, hence enhancing generalizability.

Core psychiatric issues such as comorbidity (McGrath et al., 2020) and unpredictable progression of mental disorders (Quinlan et al., 2020) hinder timely diagnosis and precision treatment. The challenge entails relatively low signal intensity specific to group-level psychological/psychiatric disorders in contrast to between subject variances. It is difficult to progress without consistent and valid data as the foundation. To reach that goal requires confronting the heterogeneity inherent in synthesizing small-scale data from diverse sources (Yan et al., 2019). The bias and variances among different scanners, selection standards and samples result in inconsistencies among sites, nonlinearly expanded along with the number of sources. Solutions responding to this need emerge especially in the intersection of medical imaging and artificial intelligence (Dinsdale et al., 2021; Guan et al., 2021; Zhong et al., 2020). However, only a few of these methods were designed for or have experimented with R-fMRI data. Moreover, few methods have thoroughly tackled efficiency, identifiability, test-retest reliability and replicability, probably due to a lack of appropriate datasets designed for such purposes. Our work fills this gap and supports the conclusion that a nonparametric algorithm fits the above requirements. SMA was inspired by causal inference. It is empowered by a flexible and free application structure, can be easily extended and modified to deal with many kinds of situations. Still, freedom has a price. The price of this flexibility is exactly what data science requires. We must dig into our data to learn its distribution, dimensions, relationships among factors, and population where it was sampled as the priors that will guide subsequent analyses. For all these reasons, the SMA method will soon be implemented in Data Preprocessing & Analysis for Brain Imaging (DPABI) (Yan et al., 2016) for convenient application in the R-fMRI field.

We note some limitations. Our analyses occurred in volume space, while studies have shown surface-based approaches enable more precise spatial localization (Coalson et al., 2018) and decrease signal contamination (Brodoehl et al., 2020). Our findings should be further validated with surface-based preprocessing methods. Moreover, guidelines on choosing the target site for optimizing SMA are preliminary. Our formula provides a preliminary mechanism for filtering target sites which should be further tested and modified as needed to enhance its generalizability.

In future, it would be preferable to establish a thorough and instructive processing standard for implementing SMA. We welcome and advocate the design and collection of large-scale test-retest datasets consisting of traveling subjects that would include more recorded variables for verification and increasing improvements. State-of-the-art techniques aimed at harmonizing R-fMRI should be compared to SMA on identifiability, test-retest reliability and generalizability as well as on applicability. For a complicated structured neural network such as ICVAE, a heavy workload is transferred to representative data accumulation and experiments to testify its generalizability. Making codes open and providing a clear instruction manual will advance the translation and application of new AI techniques regardless of users’ professional backgrounds.

## Conclusion

In this study, we selected site-harmonization approaches along the statistical theory spectrum and systematically compared them in validity, test-retest reliability, and generalizability on multiple datasets. SMA is for now the best choice for site harmonization, performing well over all perspectives. We offer our application recommendations for SMA based on experiments and we also give suggestions for future studies to harmonize site effects in brain imaging big data.

## Credit authorship contribution statement

**Yu-Wei Wang** 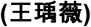: Conceptualization, Methodology, Software, Formal analysis, Investigation, Data Curation, Writing - Original Draft, Writing - Review & Editing, Visualization, Funding acquisition. **Xiao Chen** 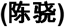: Investigation. **Chao-Gan Yan** 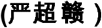: Conceptualization, Methodology, Software, Writing - Original Draft, Writing - Review & Editing, Supervision, Project administration, Funding acquisition.

## ACKNOWLEDGEMENTS

We thank Prof. Francisco Xavier Castellanos for his comments and edits. This work was supported by the Sci-Tech Innovation 2030 - Major Project of Brain Science and Brain-inspired Intelligence Technology (grant number: 2021ZD0200600), National Key R&D Program of China (grant number: 2017YFC1309902), the National Natural Science Foundation of China (grant numbers: 82122035, 81671774, 81630031), the 13th Five-year Informatization Plan of Chinese Academy of Sciences (grant number: XXH13505), the Key Research Program of the Chinese Academy of Sciences (grant NO. ZDBS-SSW-JSC006), Beijing Nova Program of Science and Technology (grant number: Z191100001119104), and the Scientific Foundation of Institute of Psychology, Chinese Academy of Sciences (grant number: E2CX4425YZ).

