## SupplementaryTables for "Comprehensive evaluation of harmonization on functional brain imaging for multisite data-fusion"

Supplementary Table 1. Dice coefficients for intrinsic brain metrics between two sessions of CoRR after FDR correction with head motion (mean FD Jenkinson), sex and age as covariates. The numbers below percentage represents the number of voxels who are significant in both sessions (Sig.=2).

|  |  |  |  | FDR | | | | |  |  |  |
| --- | --- | --- | --- | --- | --- | --- | --- | --- | --- | --- | --- |
|  | ReHo | | ALFF | | | fALFF | | DC | | FC | |
| Raw | 64% | 1683 | 71% | | 7889 | 42% | 378 | 52% | 972 | 10% | 2 |
| Reg | 66% | 1617 | 62% | | 4613 | 41% | 541 | 51% | 1008 | 15% | 2 |
| Adj | 66% | 2395 | 65% | | 5969 | 43% | 1018 | 56% | 1734 | 19% | 6 |
| LMM | 65% | 2147 | 66% | | 6222 | 42% | 714 | 53% | 1373 | 22% | 6 |
| ComBat(adj+para) | 67% | 2407 | 69% | | 7140 | 43% | 952 | 56% | 1708 | 28% | 9 |
| ComBat(adj+nonpara) | 66% | 2363 | 69% | | 7276 | 44% | 976 | 56% | 1765 | 26% | 9 |
| ComBat(unadj+para) | 67% | 1930 | 68% | | 6918 | 42% | 588 | 53% | 1121 | 29% | 7 |
| ComBat(unadj+nonpara) | 67% | 1938 | 69% | | 6973 | 42% | 584 | 53% | 1128 | 29% | 7 |
| SMA | 68% | 2786 | 67% | | 7157 | 45% | 1304 | 55% | 1852 | 24% | 14 |
| ICVAE | 52% | 2252 | 74% | | 13764 | 47% | 1291 | 45% | 1543 | NaN | 0 |

Supplementary Table 2. Dice coefficients for CORR session1, 2 and FCP1/2 (Dice31, Dice32) with 587 subjects without scanner as regressor.

| **GRF**  **corrected** | ReHo | | | | ALFF | | | | fALFF | | | | | DC | | | |
| --- | --- | --- | --- | --- | --- | --- | --- | --- | --- | --- | --- | --- | --- | --- | --- | --- | --- |
|  | Dice31 | | Dice32 | | Dice31 | | Dice32 | | Dice31 | | | Dice32 | | Dice31 | | Dice32 | |
| Raw | 14%  [131] | | | | 2%  [43] | | | | 18%  [608] | | | | | 6%  [66] | | | |
| **Empirical Bayesian ComBat** | | | | | | | | | | | | | | | | | |
| adj+para | 8%  [63] | | 7%  [59] | | 14%  [148] | | 13%  [127] | | 13%  [234] | | | 16%  [266] | | 9%  [43] | | 9%  [40] | |
| adj+nonpara | 8%  [65] | | 7%  [62] | | 13%  [135] | | 12%  [119] | | 12%  [222] | | | 2%  [240] | | 10%  [45] | | 8%  [35] | |
| **Maximum mean distance** | | | | | | | | | | | | | | | | | |
| SMA | 12%  [134] | | 11%  [130] | | 12%  [211] | | 12%  [209] | | 19%  [466] | | | 20%  [512] | | 10%  [78] | | 9%  [70] | |
| **FDR**  **corrected** | ReHo | | | ALFF | | | | fALFF | | | DC | | | | FC | | |
|  | Dice31 | Dice32 | | Dice31 | | Dice32 | | Dice31 | | Dice32 | Dice31 | | Dice32 | | Dice31 | | Dice32 |
| Raw | 17%  [426] | | | 10%  [470] | | | | 36%  [3144] | | | 6%  [185] | | | | 1%  [1] | | |
| **Empirical Bayesian ComBat** | | | | | | | | | | | | | | |  | |  |
| adj+para | 16%  [320] | 17%  [426] | | 13%  [311] | | 13%  [285] | | 29%  [1649] | | 29%  [1654] | 10.0%  [111] | | 9.0%  [85] | | 0%  [0] | | 1%  [2] |
| adj+nonpara | 15%  [331] | 15%  [310] | | 13%  [285] | | 12%  [268] | | 30%  [1758] | | 30%  [1783] | 10.0%  [120] | | 9.0%  [92] | | 0%  [0] | | 1%  [2] |
| **Maximum mean distance** | | | | | | | | | | | | | | |  | |  |
| SMA | 18%  [578] | 18%  [555] | | 19%  [801] | | 19%  [775] | | 35%  [2462] | | 37%  [2643] | 17%  [401] | | 17%  [401] | | 0%  [0] | | 4%  [1] |

Supplementary Table 3. Clustering accuracy of target site choice experiments on TSP-3.

| TST DATASET:  Clustering Accuracy | zReHo | | zALFF | | | zfALFF | | | zDC | | | FC-142ROI | |
| --- | --- | --- | --- | --- | --- | --- | --- | --- | --- | --- | --- | --- | --- |
|  | Euclidean | Pearson | Euclidean | Pearson | Euclidean | | Pearson | Euclidean | | Pearson | Euclidean | | Pearson |
| IPCAS-GE | 85% | 88% | 85% | 95% | 44% | | 56% | 63% | | 59% | 66% | | 90% |
| PKU-GE | 88% | 93% | 83% | 93% | 59% | | 63% | 66% | | 71% | 68% | | 93% |
| PKU-SIEMENS | 93% | 88% | 93% | 93% | 68% | | 68% | 66% | | 71% | 66% | | 90% |

Supplementary Table 4. Stability of target site choice experiments on CoRR. We execute SMA harmonization and pointing the same target site for each session. Then we calculate the test-retest reliability between the harmonized within one session targeted different sites and average the Dice results by target site.

| **Correction Method** | GRF | | | | | FDR | | | | |
| --- | --- | --- | --- | --- | --- | --- | --- | --- | --- | --- |
| **Target Site** | BNU | IACAS | JHNU | MRN | SWU | BNU | IACAS | JHNU | MRN | SWU |
| ReHo | 85% | 79% | 81% | 83% | 84% | 86% | 80% | 83% | 84% | 85% |
| ALFF | 88% | 83% | 84% | 86% | 87% | 88% | 85% | 84% | 87% | 88% |
| fALFF | 87% | 85% | 83% | 84% | 87% | 90% | 88% | 87% | 88% | 90% |
| DC | 82% | 79% | 80% | 80% | 79% | 84% | 81% | 82% | 84% | 83% |
| FC |  |  |  |  |  | 75% | 61% | 65% | 71% | 73% |
