## SupplementaryFigures for "Comprehensive evaluation of harmonization on functional brain imaging for multisite data-fusion"

Supplementary Figure 1. **ICVAE model structure and parameters.** We use the same network structure for both TSP-3 and CoRR, only change the dimensions of intermediate z and output, in accordance to their site number (3 for TSP-3 and 6 for CoRR).

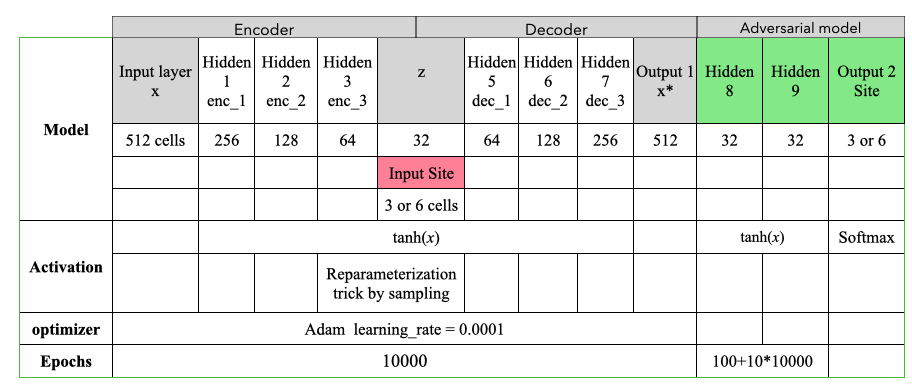

Supplementary Figure 2. **Overlapped sex statistical results on brain for GRF corrected** **fALFF, ReHo and DC.**
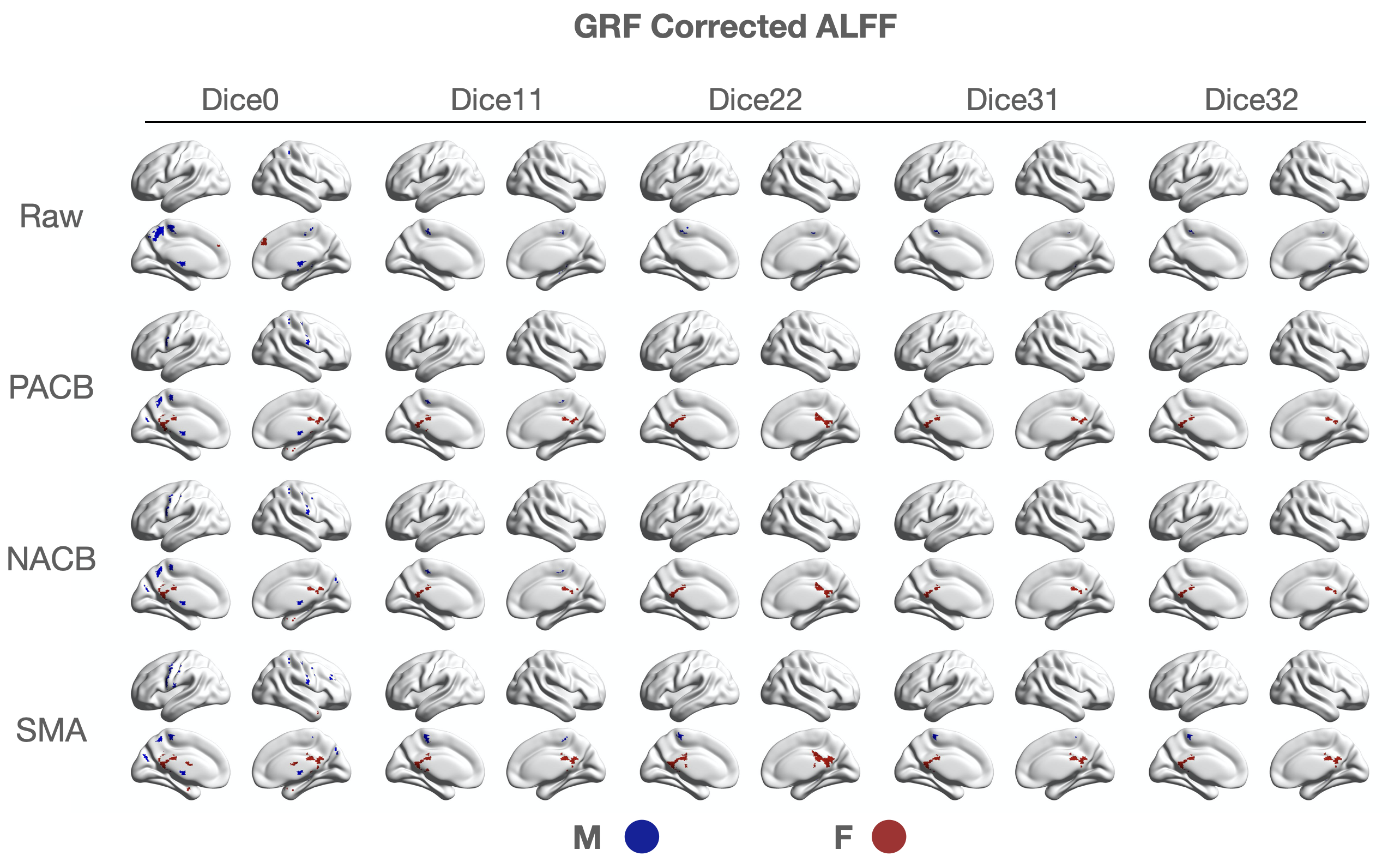

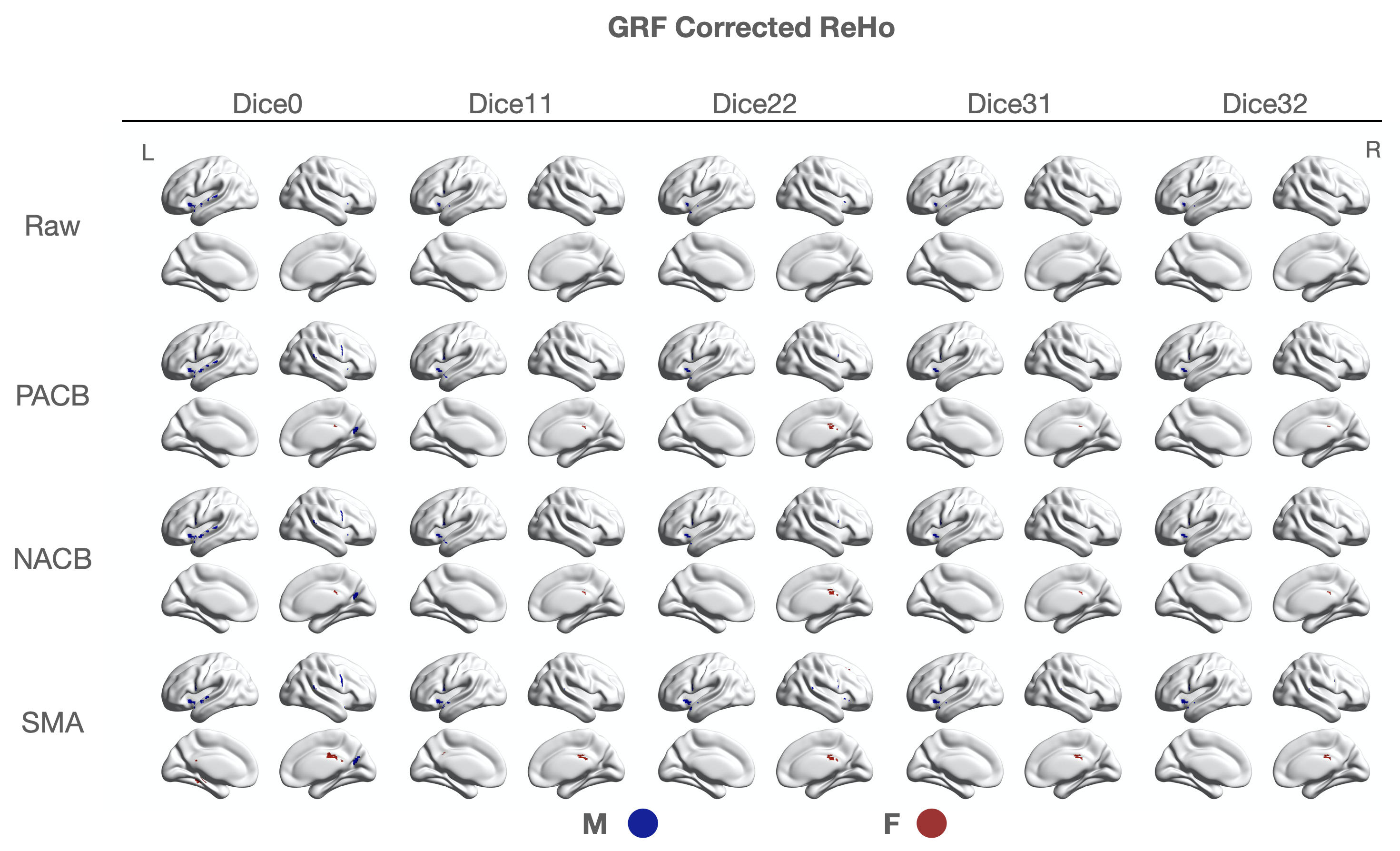

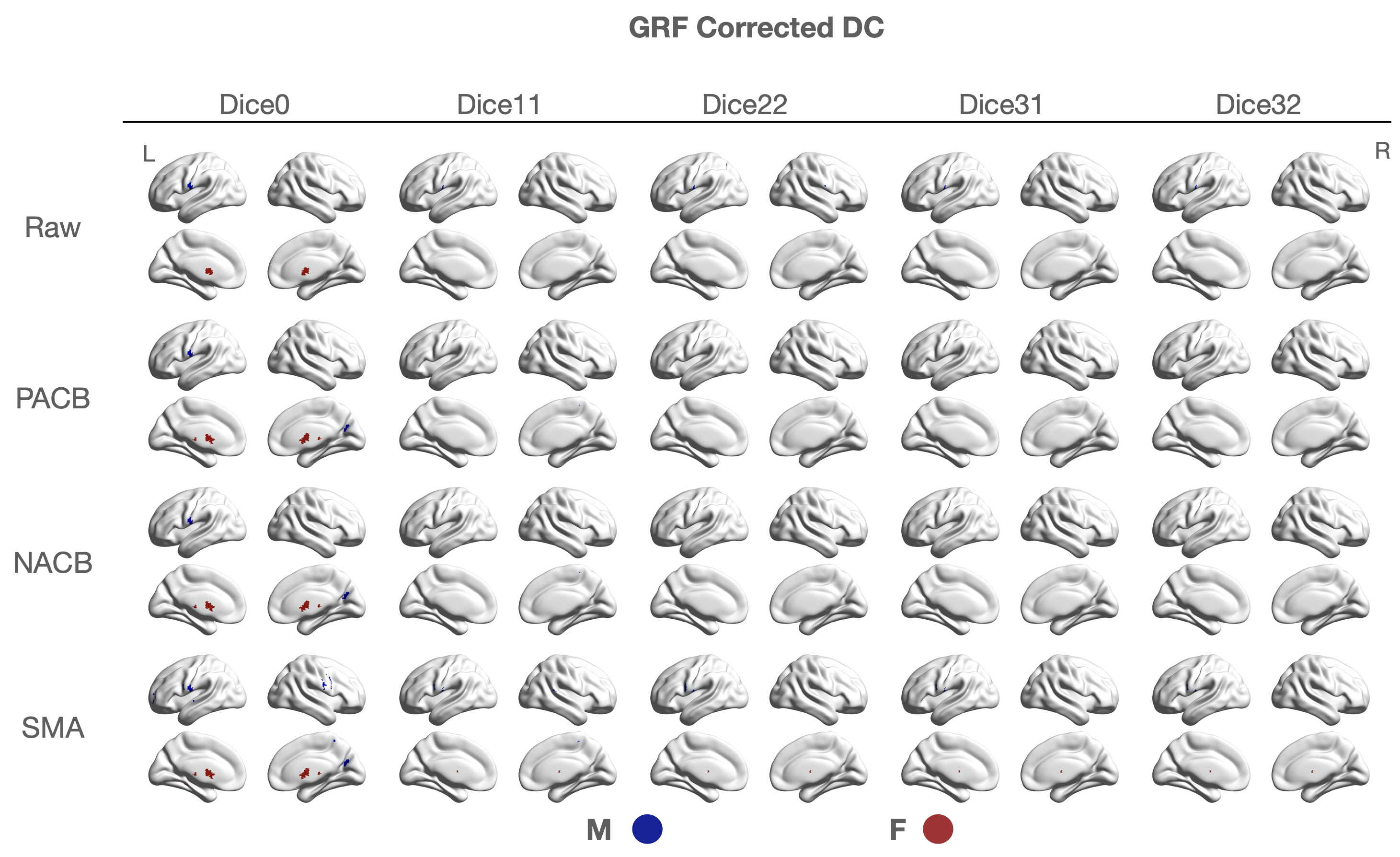

Supplementary Figure 3. **Site choice experiments**. CoRR: Test-retest reliability and stability when choosing different sites as target. FDR corrected. The x-axis is arranged in order of site selection formula, left to right best to worst. Color and shape of points distinguish R-fMRI metrics.

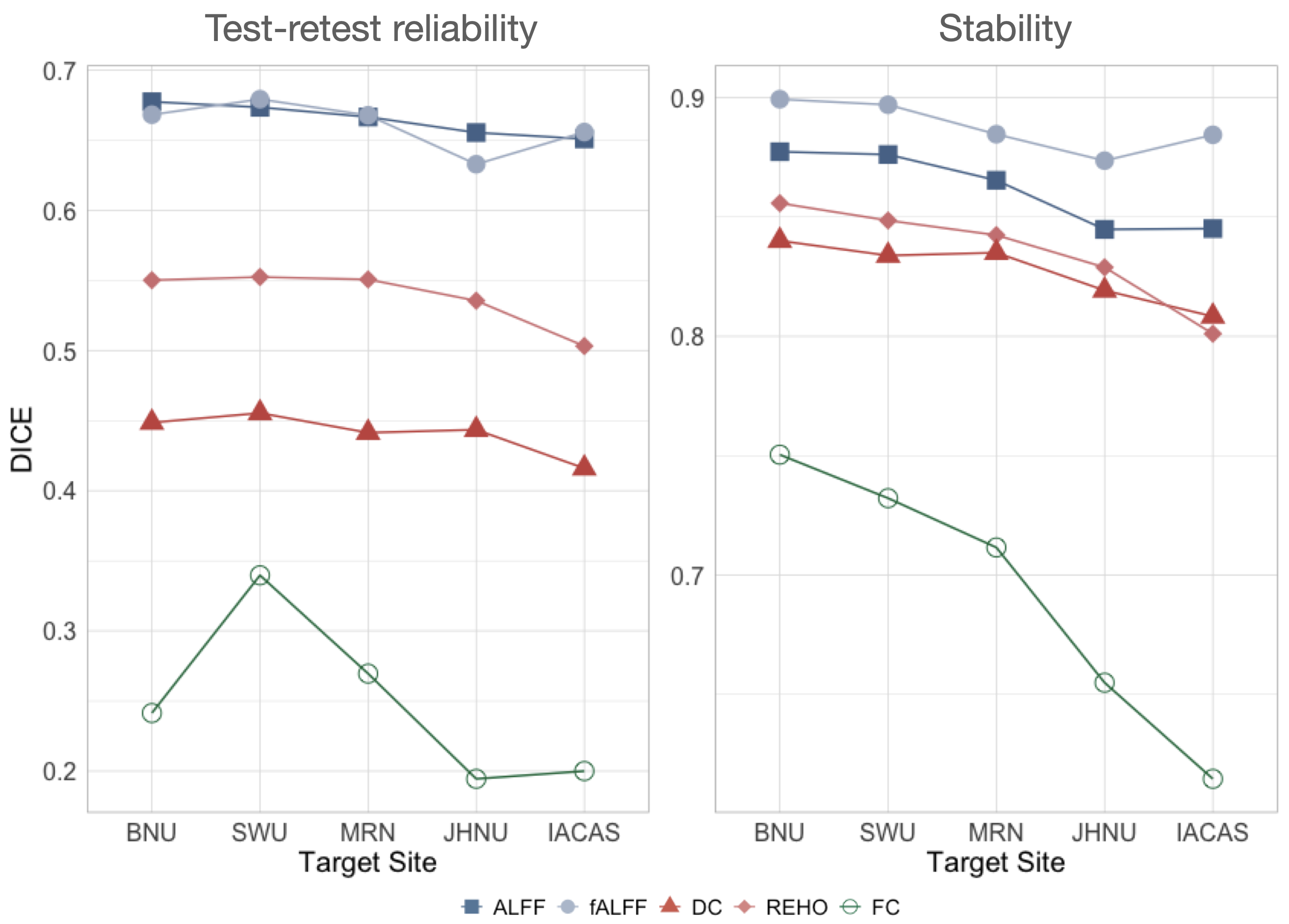
